# TALEN-induced contraction of CTG trinucleotide repeats in myotonic dystrophy type 1 cells

**DOI:** 10.1101/2023.10.14.562330

**Authors:** Laureline Bétemps, Stéphane Descorps-Declère, Olivia Frenoy, Lucie Poggi, Valentine Mosbach, Stéphanie Tomé, David Viterbo, Arnaud Klein, Laurence Ma, Sonia Lameiras, Thomas Cokelaer, Marc Monot, Bruno Dumas, Geneviève Gourdon, Denis Furling, Guy-Franck Richard

## Abstract

Trinucleotide repeat expansions are the cause of two dozen neurodegenerative and developmental disorders. One of these, myotonic dystrophy type 1 (Steinert disease, or DM1) is due to the expansion of a CTG triplet in the 3’ UTR of the *DMPK* gene. We used highly specific DNA endonucleases to induce a double-strand break in the repeat tract to contract it below pathological length. Expression of a TALE Nuclease (TALEN) in human DM1 cells induced moderate CTG repeat contractions in 27% of the clones analyzed. These clones exhibited large internal deletions within the TALEN, occurring by homologous recombination between internal TALE repeats, inactivating the nuclease, and explaining its reduced efficacy. Taking advantage of the degeneracy of the genetic code, we recoded the TALEN sequence, to decrease internal redundancy and optimize codon usage. The new recoded TALEN showed increased efficacy in DM1 cells, with 68% of clones exhibiting a moderate to large contraction of the CTG repeat tract. In contrast, *Staphylococcus aureus* Cas9 (*Sa*Cas9) was unable to contract the CTG repeat tract. In parallel, we completely sequenced to very high coverage the DM1 genome using the PacBio technology. Several clones in which the TALEN was induced were also totally sequenced. In some of them, length changes of other long CTG repeats were detected, possibly corresponding to off-target effects, all of them in introns or intergenic regions. Repeat contractions were never associated with recombination of flanking markers, suggesting that contractions most probably occur by an intra-allelic mechanism such as single-strand annealing. TALENs should now be considered as a promising gene therapy approach, not only for DM1 but also for many other microsatellite expansion disorders.

## Introduction

Microsatellite expansions are responsible for over two dozen human disorders, including Huntington’s disease, myotonic dystrophy type 1 (DM1 or Steinert disease), Friedreich’s ataxia, fragile-X syndrome and various types of ataxias as well as severe developmental pathologies (reviewed in Orr, 2001; Orr & Zoghbi, 2007). Among these, DM1 arises from the expansion of the CTG triplet repeat within the 3’ UTR of the *DMPK* gene, on chromosome 19 (Brook *et al* , 1992). DM1 is characterized as a dominant autosomal multisystemic disorder, encompassing myotonia, progressive muscle weakness and wasting, cataracts, cardiac conduction defects and cognitive dysfunction. These symptoms consistently coincide with an expansion of the *DMPK* repeat tract. In individuals without the disease, the CTG repeat length typically falls within the range of 5 to 37 triplets, whereas patients exhibit lengths ranging from 50 to 4,000 CTG. However, even among the different forms of the disease, there is substantial variability in the length of the repeat tract. For instance, the adult form of the disease can feature repeat expansions ranging from 250 to 750 CTG triplets, while the infantile form may have expansions ranging from 500 to over 1000 triplets (De Antonio *et al*, 2016). Consequently, for a given repeat length, predicting the severity of the symptoms is challenging. A patient with 500 CTG triplets could present the adult, juvenile, of infantile form of the disease, each associated with various degrees of symptom severity. This incomplete penetrance implies the involvement of additional genetic factors, beyond the *DMPK* CTG expansion, in determining DM1 severity.

At the cellular level, patient’s cells exhibit abnormal nuclear aggregates of mRNA carrying expanded CUG repeats. These CUGexp-mRNA sequester the MBNL RNA binding proteins, leading to dysregulations in RNA metabolism, such as alternative splicing defects and multisystemic deficiencies that ultimately contribute to the pathology. At the molecular level, it is the expression of pathological CUGexp-mRNA that triggers the disease. Therefore, several therapeutic approaches have been developed to target this expanded transcript, either directly or indirectly. Strategies such as inhibiting or reducing *DMPK* expression using small molecules or antisense oligonucleotides, have been explored, with several of them entering clinical trials (Pascual-Gilabert *et al* , 2021). In another approach, the expanded CUG mRNA was targeted by a mutated version of *Streptococcus pyogenes* Cas9 (*Sp*Cas9). The presence of the deficient nuclease was sufficient to decrease the amount of toxic mRNA in cell cultures and suppress RNA foci and some splicing defects (Batra *et al*, 2017).

Another therapeutic approach involved targeting the CUGexp-mRNA to prevent its interaction with the MBLN1 protein, using antisense oligonucleotides (reviewed in Izzo *et al*, 2022). More recently, a gene therapy strategy using a decoy with high affinity for CUGexp has also been developed in order to disrupt the deleterious interaction between MBNL1 protein and the expanded transcript (Arandel *et al*, 2022).

Another avenue of research focuses on the development of gene therapy approaches targeting the expanded CTG trinucleotide repeat at the DNA level, using highly specific DNA endonucleases such as Zinc-Finger nucleases (ZFN), TALE nucleases (TALEN) or the large CRISPR-Cas family (Mosbach *et al*, 2018a). Two main strategies have been pursued: one aims to contract the expanded CTG repeat by induction of a double-strand break (DSB) within the repeat tract, while the other seeks to delete it by inducing DSBs upstream and/or downstream of the repeat. The latter strategy has shown some success in DM1 cells and transgenic mice using either a modified version of the *Streptococcus pyogenes* Cas9 (e*Sp*Cas9) or the wild-type *Staphylococcus aureus* Cas9 (*Sa*Cas9) (Provenzano *et al* , 2017; Lo Scrudato *et al* , 2019; Dastidar *et al*, 2018), although it sometimes results in large deletions around the repeat tract (Agtmaal *et al* , 2017). Similar deletions have been observed in a yeast model containing an expanded CTG repeat from a DM1 patient, where *Sp*Cas9 was used to induce a DSB within the repeat tract (Mosbach *et al* , 2020). An alternative strategy proposes to induce a DSB within the expanded CTG trinucleotide repeat in order to contract it, hopefully alleviating the symptoms of the pathology (Richard, 2015). A ZFN targeting an expanded CTG repeat was expressed in HeLa cells and triggered efficient replication-dependent shortening of the repeat tract (Liu *et al* , 2010). The *Sp*Cas9 nickase was also shown to contract the expanded CTG triplet repeat in an HEK293 cell model, although with a low but detectable efficacy (Cinesi *et al*, 2016). A TALEN was employed for the same purpose and demonstrated to induce specific contractions of an expanded CTG repeat from a DM1 patient in a *Saccharomyces cerevisiae* yeast model, achieving 99% efficacy (Richard *et al* , 2014). Efficient DSB resection by the Mre11-Rad50 complex as well as the presence of the associated Sae2 protein were necessary to shorten the repeat tract (Mosbach *et al*, 2018b).

Here, we induced both TALEN and *Sa*Cas9 in fibroblasts derived from transgenic DM1 as well as immortalized DM1 fibroblasts. Notably, *Sa*Cas9 exhibited diminished or negligible efficacy in contracting the CTG repeat tract when compared to TALEN. We observed frequent internal deletions occurring in both arms of the TALEN following transduction in human cells. To address this issue, we recoded the TALEN using the degeneracy of the genetic code, thereby reducing motif similarity and adapting it to the *Homo sapiens* codon usage preference. This newly recoded TALEN exhibited superior efficacy in contracting the CTG repeat tract compared to the wild-type nuclease. Subsequently, we conducted PacBio whole-genome sequencing and uncovered contractions in other long CTG repeats within the human genome. These findings support the presence of off-target mutations induced by the TALEN in other long microsatellites.

## Results

### TALEN expression in mammalian cells

To ensure precise targeting of the DMPK locus, the TALEN was engineered in such a way that the left arm binds the CTG repeat tract and the right arm was designed to recognize the boundary between the repeated and non-repeated sequence (Figure 1A). In our previous work, we have reported that breaks were induced approximately 3-4 triplets before the end of the repeat tract (Mosbach *et al* , 2018b). Both TALEN arms were cloned in a plasmid (pMOV1) flanking a GFP gene, to prevent recombination between TALEN arms following transfection (Figure 1A) (Boch *et al*, 2009). Fibroblasts derived from a hemizygous transgenic mice carrying the human *DMPK* gene with 600 CTG triplets were transfected with pMOV1. GFP-positive cells were sorted, and total genomic DNA was extracted for repeat length analysis using PCR. Notably, the amplification of the repeat tract revealed that the expanded allele had contracted to less than approximately 30 CTG triplets, while control cells showed no significant length difference (Figure 1B).

**Figure 1:**
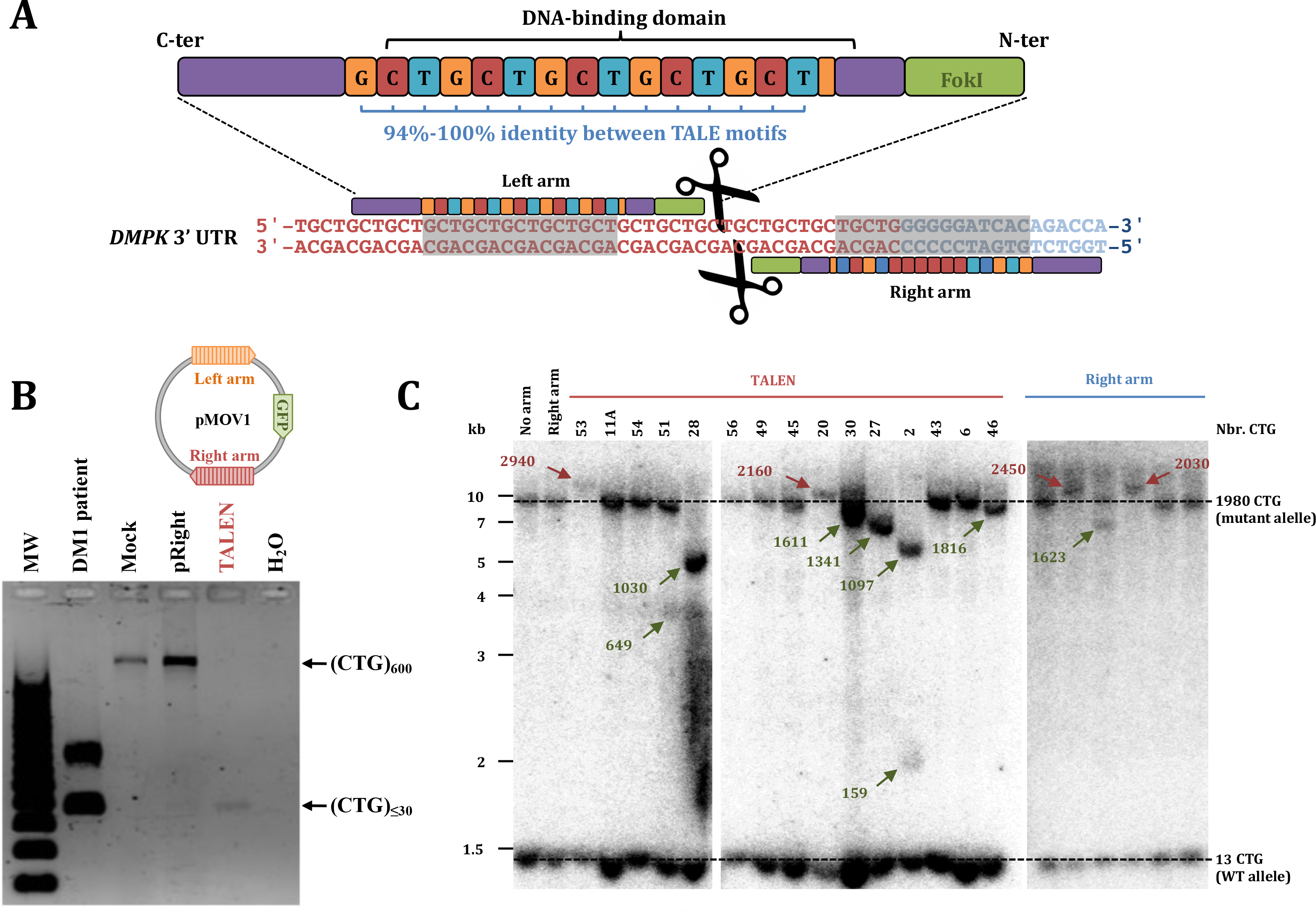
TALEN design and expression in mice and human cells. A: TALEN design, showing the respective binding sites of each arm at the 3’ end of the *DMPK* gene (Richard *et al* , 2014). **B:** TALEN expression in mice fibroblasts containing 600 CTG triplets. pRight: transfection with the right arm only. TALEN: transfection with both arms (pMOV1). The CTG repeat was amplified with primers ST300f and ST300r. C: TALEN expression in ASA cells. The Southern blot shows independent clones in which the right arm alone or both arms (TALEN) were transduced. Red arrows point to expansions, green arrows to contractions. Repeat lengths are shown in triplets, as determined by measurements on the blot.

Each TALEN arm was subsequently inserted into two lentivirus backbones bearing different selection markers. Immortalized human DM1 fibroblasts, referred to as ASA cells (Arandel *et al*, 2017), underwent a two-step transduction process. First, they were transduced with the lentivirus carrying the right arm and selected for neomycin resistance. Subsequently, they were transduced with the lentivirus carrying the left arm and selected for hygromycin resistance. Cells that successfully incorporated both arms were expanded clonally, and genomic DNA was extracted to assess CTG repeat length by Southern blotting. In this particular case, PCR could not be used due to the presence of an expanded *DMPK* allele with ∼2,000 triplets. Cells with only one TALEN arm exhibited a low level of repeat instability, with one expansion and two contractions identified among 28 alleles analyzed. Cells in which both arms were expressed showed two expansions and 10 contractions among 37 alleles analyzed. The slight increase in contractions when both arms are expressed did not reach statistical significance. Additionally, most of the observed contractions were partial, with some clones showing more than one contracted allele (Figure 1C). Note that we focused only on the expanded allele, as detecting length changes in the smaller wild-type allele at this scale was not feasible. This series of experiments indicated that the wild-type TALEN demonstrated high efficiency in contracting a (CTG)_600_ repeat in mice fibroblasts but displayed reduced efficacy in contracting a (CTG)_2000_ expansion in human DM1 cells.

### Analysis of CTG trinucleotide repeats by PacBio sequencing of ASA cells

A crucial concern with gene editing is the precision of genetic modifications. Off-target mutations are a common occurrence when utilizing *Sp*Cas9 and their prediction is not always feasible through computational tools (Tsai *et al* , 2015). In the human reference genome (GRCh38), it is estimated that there are at least 10,000 CTG trinucleotide repeats (or their complementary sequence CAG) (International Human Genome Sequencing Consortium, 2001). However, this figure is likely an underestimate, particularly because sequencing very long repeats poses challenges when employing short read technologies. A more comprehensive sequencing approach, incorporating a combination of short, long and very long reads, as well as Hi-C and optical mapping, has recently led to the release of the first human genome sequence from telomere to telomere (T2T), including all centromeres that were known to contain very large repeated elements. Initial analysis of this genome sequence has suggested the presence of potentially twice as many microsatellites as previously identified, likely encompassing very long ones that were previously beyond the reach of sequencing techniques (Nurk *et al* , 2022). Consequently, we made the decision to employ long-read sequencing as a means to evaluate possible off-target effects of the TALEN.

The ASA genome underwent sequencing using the PacBio technology. Subreads obtained from four Sequel II SMARTCells covered a total of 1,381 nucleotides. When converted to HiFi reads, 82.5 Gb of high-quality sequence were obtained, representing a 13X coverage of the human haploid genome. These reads were employed for the primary assembly, yielding 371 contigs (N50: 52.9 Mb), among which 69 are sufficient to cover 90% of the genome (Supplemental Table S1). Notably, this marks the first instance, to the best of our knowledge, of a DM1 genome sequenced and phased with long read technology. To comprehensively identify all potential CTG trinucleotide repeats, both perfect and imperfect, we developed an in-house analysis pipeline, named OSTINATO (Descorps-Declère and Richard, unpublished). This program was executed on the ASA3-p11 reference genome and led to the detection of 8 imperfect and two perfect CTG trinucleotide repeats spanning more than 48 triplets, approximately the size at which the *DMPK* CTG allele becomes pathogenic. Each of these long CTG repeats was given a unique identifier related to the cytogenetic band to which it belongs. They range in size from 48 to 278 CTG triplets (Table 1).

For comparative purposes, we assessed the performance of our software against the established gold standard TRF program, which is well-recognized and widely used for tandem repeat analysis (Benson, 1999). Remarkably, TRF did not detect 2 out of the 8 imperfect CTG repeats identified by OSTINATO.

All reads covering these 10 repeated regions were extracted and visually inspected. Repeat lengths were not strictly identical among all reads. This may be due to somatic mosaicism or to reduced sequencing accuracy within these long repeated regions. The median length of each allele was therefore used for subsequent analyses. Each identified repeat displayed one or two distinct median repeat lengths, indicative of homozygous or heterozygous loci. The two alleles detected at the DMPK locus contain respectively 13 and 1806 CTG triplets, in the reference ASA3-p11 genome (Table 1). The complete list of HiFi reads covering the *DMPK* locus for each clone, with the length of CTG repeat tracts is given in Table 2.

It is noteworthy that the imperfect CTG repeat lengths identified by TRF were significantly longer that those identified with OSTINATO. This discrepancy arises from the implementation of different algorithms for imperfect repeat detection. Both programs yielded consistent results for the two perfect CTG repeats (17q21_2 and 19q13). However, TRF was unable to detect the short wild-type DMPK allele given the detection threshold used (Table 1).

### For: Open Solution Tallying Iterative Nucleotides Aligned in Tandem Order

Subsequently, ten independent ASA clones in which the TALEN was transduced were sequenced with long reads, and their genome was analyzed using the same pipeline. A comprehensive list of all sequenced clones can be found in Supplemental Table S2 and genome coverages are shown in Supplemental Table S3.

Due to low sequence quality in repeated regions, CTG repeat lengths were not strictly identical among all reads covering the same genome. For that reason, we considered that a repeat length was statistically different from the reference when its median length was more than two standard deviations away from the reference median length. These occurences were counted as contractions of the CTG repeat (or in one case, expansion). In the ten clones in which both TALEN arms were transduced, all but one length alterations at the *DMPK* locus were contractions. One expansion was detected in the ASA51 clone but in only one read. Note that both alleles were not detected in all clones due to partial sequence coverage (ND in Table 1). The mutant allele is slightly underrepresented as compared to the wild-type allele, because reads that ended up within the expanded CTG triplet were discarded since their length could not be precisely determined.

When feasible, the expanded allele lengths measured via Southern blots were compared with those obtained through long read sequencing. This comparison revealed a strong correlation between the two methods (Supplemental Figure S1).

These findings confirmed the observation obtained by Southern blotting, indicating that TALEN expression predominantly led to contractions of the *DMPK* CTG repeat, albeit often partial. In one instance, a limited expansion was detected. However, this expansion should be carefully taken into account since it was found in only one read.

Moderate contractions were also observed in eight of the nine other long CTG repeats, in several clones. These may be interpreted either as somatic instability of long CTG repeats or as possible off-target DSBs induced by the nuclease (Table 1).

### Both TALEN arms carry frequent internal deletions inactivating them

The utilization of lentiviral particles to deliver the TALEN allowed us to investigate their integration sites within the ASA genome. In each clone, one to three integration sites were detected (Figure 2). At least one right arm and one left arm were successfully identified in the sequence, with the exception of ASA22C in which only the right arm could be detected. This likely stems from incomplete genome coverage in this particular clone. To our surprise, all TALEN arms identified exhibited extensive internal deletions within the DNA-binding region. These deletions varied in size, ranging from 8 TALE motifs to encompassing the entire 15 motif DNA-binding region (Figure 2A). This observation mirrored what had been previously described for TALENs expressed from lentiviruses (Holkers *et al* , 2013). These deletions strongly indicated that the TALEN activity had been inactivated by these mutations, providing a plausible explanation for the reduced effectiveness in inducing contractions. In addition, in some cases TALE motifs were present but exhibited several mutations. Examination of all reads covering the region containing the arms showed that these apparent mutations were due to low quality sequences and were therefore not real mutations (grey squares in Figure 2).

**Figure 2:**
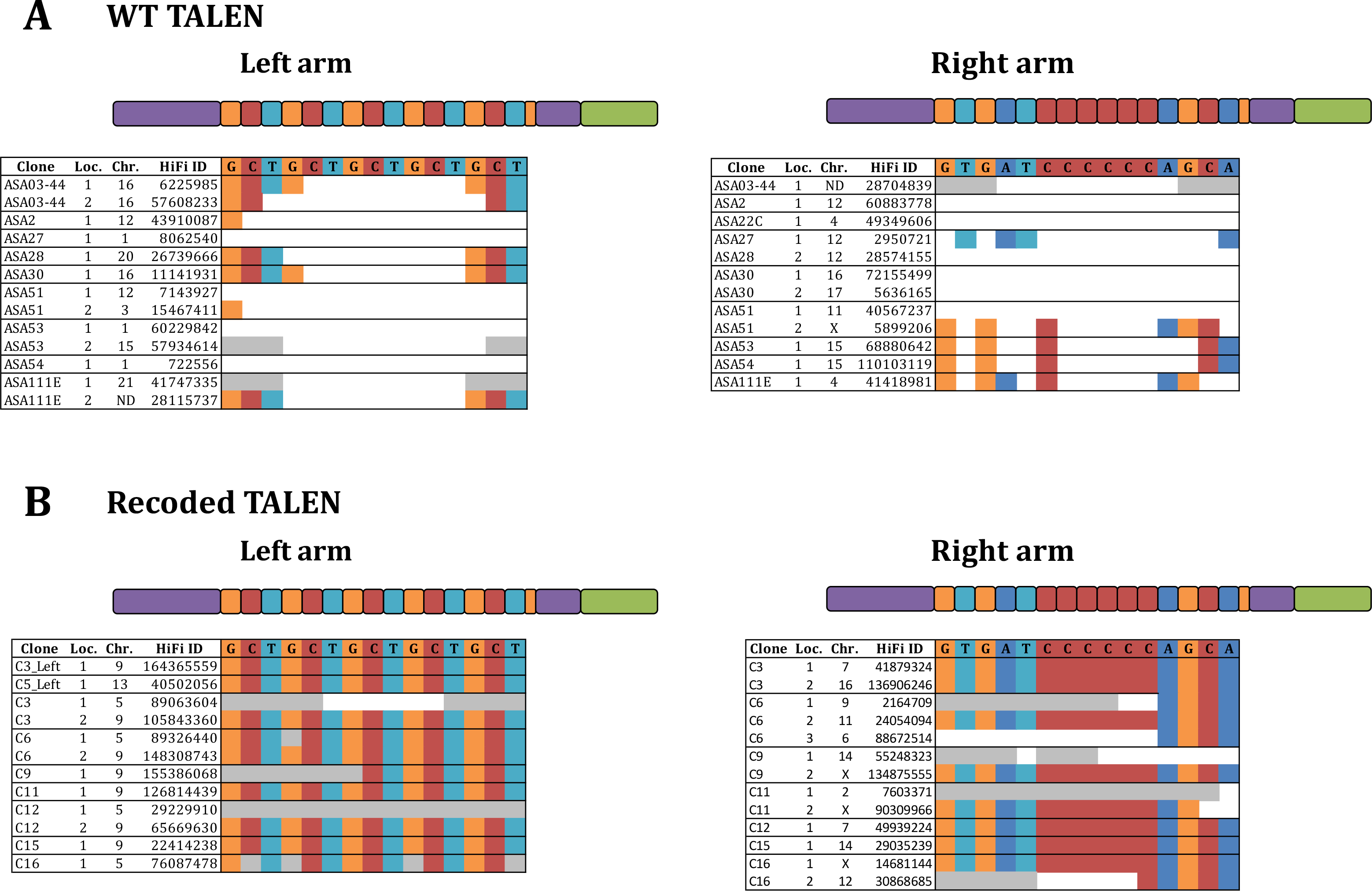
Genomic locations and integrity of TALEN arms after transduction in ASA cells. Each line corresponds to the integration of one TALEN arm in one chromosome (Chr.) at one locus in the ASA genome (Loc.). The HiFi read covering the arm is indicated (HiFi ID). Note that several reads usually span the same region but only one read is shown for each locus. Empty boxes show the extent of internal deletions in each arm. Grey boxes correspond to low quality reads. Note that for the wild-type right arm it is impossible to determine which one of the six TALE repeats binding to a cytosine is left in ASA51, ASA53, ASA54 and ASA111E. The first repeat has therefore arbitrarily been chosen. The same is true for the wild-type left arm, it is not possible to determine which ones of the TALE repeats binding to the CTG sequence were left. Flanking repeats have therefore been arbitrarily chosen. ND: the exact location of integration in the genome could not be determined because sequence quality was too low.

Pinpointing the exact timing of internal deletions occurrence during the experiment was not feasible, as all clones underwent several cell divisions before being analyzed. These deletions could potentially arise through homologous recombination between repeated motifs within each DNA-binding domain, or through template switching between internal TALE repeats during reverse transcription of the lentiviral genome, owing to the limited processivity of the reverse transcriptase (Holkers *et al*, 2013).

To address this crucial issue and enhance TALEN efficacy, we posited that reducing nucleotide identity between TALE repeats might also decrease the frequency of template switching. Consequently, we undertook a recoding approach for both TALEN arms, ensuring that contiguous motifs shared no more than 20 identical nucleotides. In addition, the sequence was optimized using the human codon index to eliminate rare codons. A prior study from the Church lab had demonstrated that recoded TALENs maintained comparable efficacies to wild-type TALENs in human cells (Yang *et al*, 2013). Using an in-house program, we generated two DNA sequences encoding the same exact TALEN protein arms, but decreasing nucleotide identity between TALE repeats from 94%-100% to 65%-78% (Supplemental Figure S2).

### An engineered recoded TALEN is more efficient at generating CTG repeat contractions

Both recoded TALEN arms (referred to as TALEN_RECODED_) were cloned into the same lentiviral vectors and transduced in ASA cells, following the same procedure. These transduced cells were clonally expanded, and total genomic DNA was extracted and analyzed by Southern blotting. The CTG repeat length prior to transduction was slightly longer than in the initial experiment with the wild-type TALEN, primarily due to several cells divisions occurring between the two experiments. It was estimated to be 2,800 triplets by Southern blotting (± 750 triplets, given resolution).

Interestingly, it was observed that repeat contractions occurred in 50% of clones that had been transduced with only the left TALEN_RECODED_ arm. This suggests that this arm, by itself, has the potential to induce CTG repeat contractions with significant frequency.

However, contractions increased to 68% when both arms were transduced. In several of these clones, contractions were nearly complete (Figure 3). In certain cases, a smear corresponding to various CTG repeat lengths was also observed. The frequency of contractions observed with this new TALEN is significantly higher than with the wild-type TALEN (Fischer exact test, p-value= 0.04).

**Figure 3:**
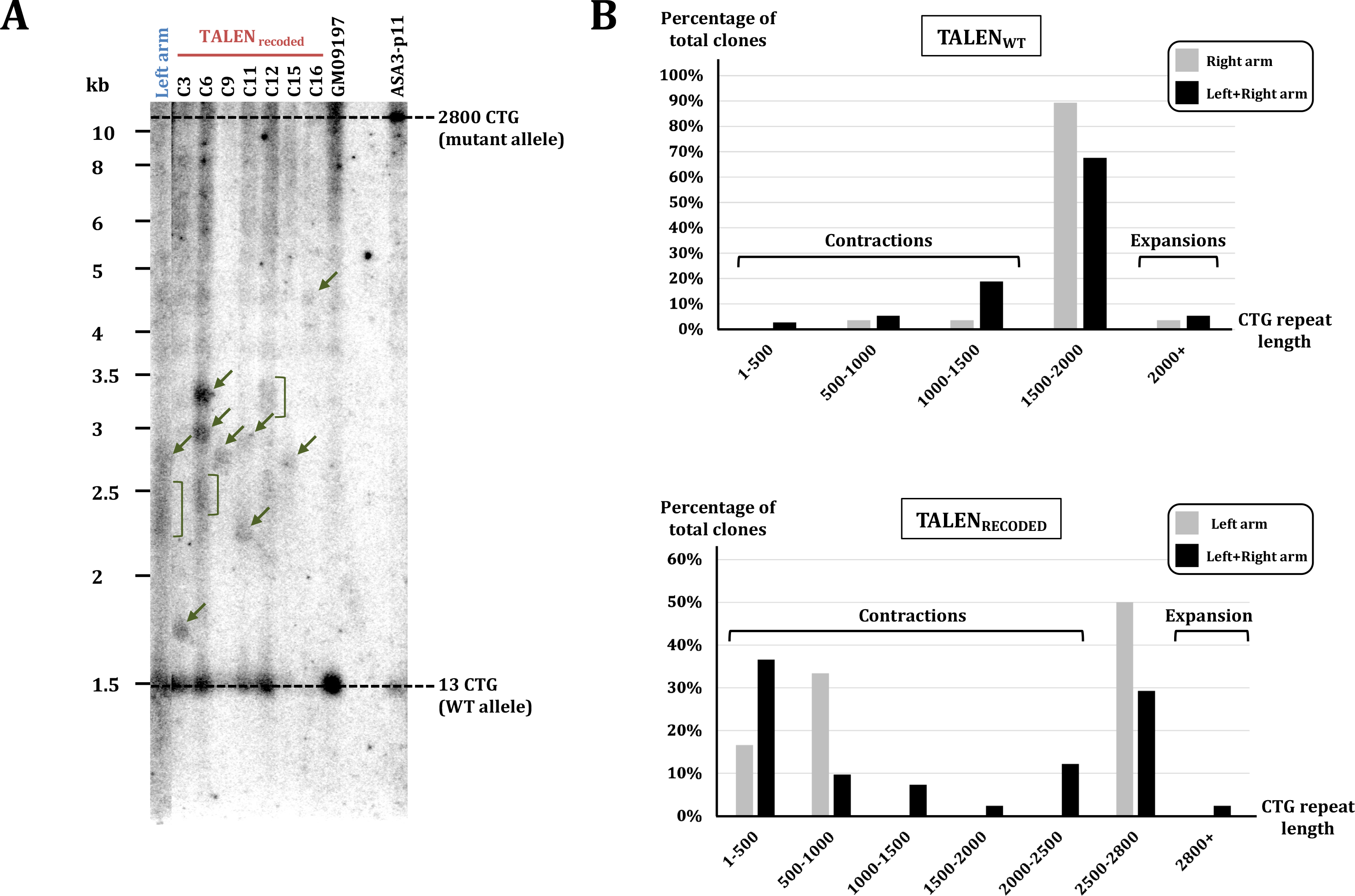
TALEN**_RECODED_** transduction in ASA cells. **A:** The Southern blot shows independent clones in which the left arm alone (Left) or both arms (TALEN_RECODED_) were transduced. GM09197: Huntington’s disease patient fibroblasts used as a control. ASA3-p11: ASA cells before transduction (reference genome). Green arrows point to contractions, green brackets to smears. **B:** length distributions of CTG repeats at the *DMPK* locus following transductions with the TALEN_WT_ (top) or the TALEN_RECODED_ (bottom).

Subsequently, nine clones transduced with the TALEN_RECODED_ were sequenced using the PacBio technology, as previously. Among them, two contained only the left arm, while the remaining seven were transduced with both TALEN_RECODED_ arms (Supplemental Table S2).

We observed that the two clones transduced solely with the left arm exhibited CTG repeat contractions at the *DMPK* locus and at the 15q23_2 locus, confirming Southern blot analysis (Table 1). This observation implies that the presence of the left arm, which binds to the repeat tract, suffices to induce CTG repeat instability. This instability may arise from the generation of double-strand breaks, single-strand breaks or by interference with the replication machinery through binding to repeat tracts. Clones in which both arms were transduced displayed more frequent contractions of the expanded *DMPK* allele, as well as alterations in the lengths of one other CTG repeat, called 15q11. This may again be interpreted as somatic mosaicism or off-target mutations. Given that these contractions are less frequent than with the wild-type TALEN (TALEN_WT_), it is more likely that they are off-target contractions. However, we cannot totally exclude that the extra contractions observed with the TALEN_WT_ come from the lower sequence quality and coverage of these clones as compared to clones induced with the TALEN_RECODED_. In total, 21 contractions were detected. No expansion was observed in the *DMPK* gene, as opposed to the wild-type TALEN. Additionally, the smears and multiple bands observed by Southern blotting for the left arm alone and for clones C6 and C12 (Figure 3) corresponded to multiple PacBio reads (Table 1). Overall, the median length of CTG repeats at the *DMPK* locus exhibited a statistically significant shift towards smaller sizes with the TALEN_RECODED_ in contrast to the wild-type TALEN (Figure 4, t-test p-value= 7.8 x 10^-12^). This shift demonstrates that the TALEN_RECODED_ was considerably more effective at contracting the CTG repeat tract than the TALEN_WT_.

**Figure 4:**
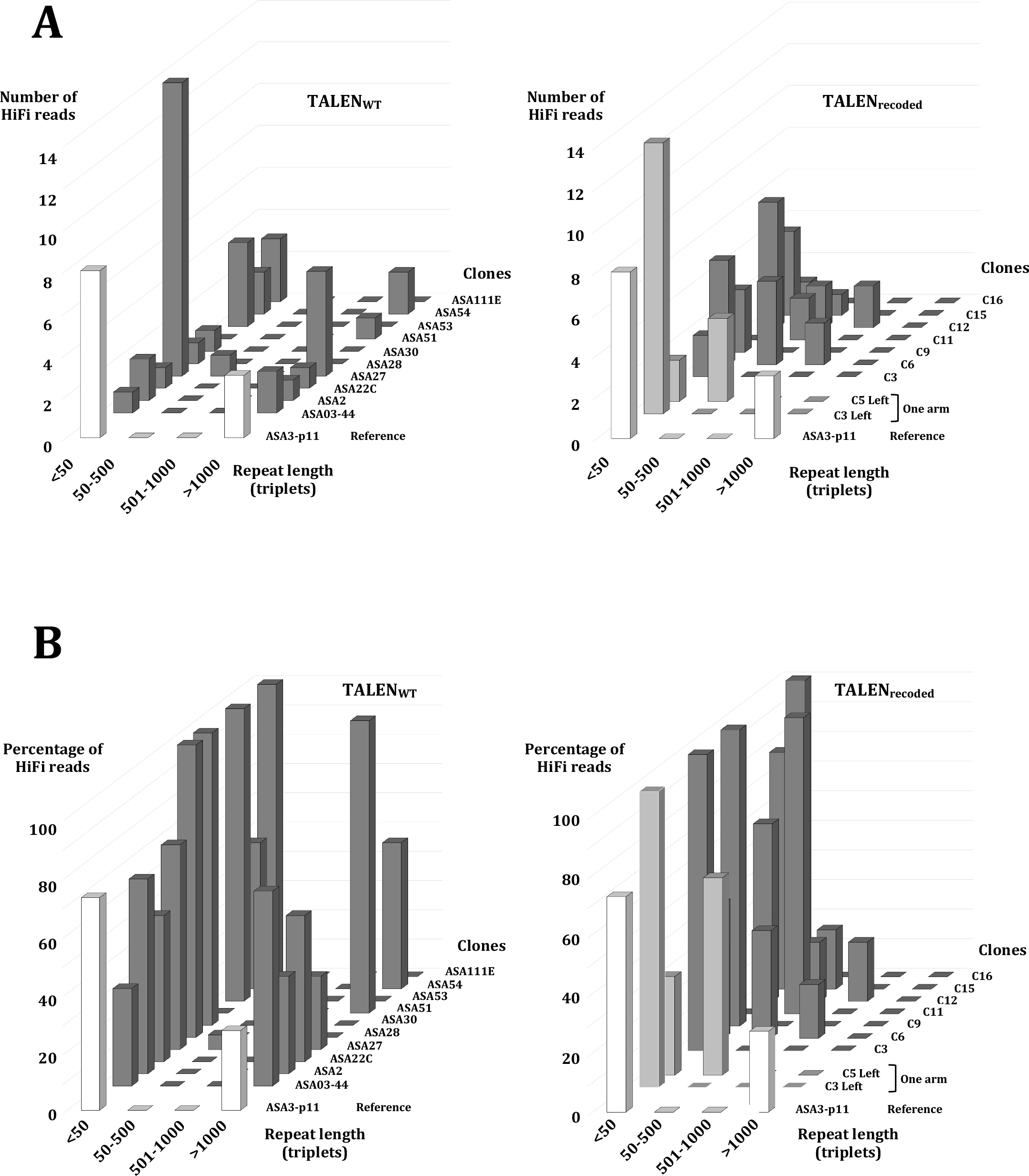
Length distributions of CTG repeats at the DMPK locus, following TALEN **_WT_** or TALEN **_RECODED_** expression. **A:** The X axis shows the length categories (in triplets), the Y axis shows the number of reads in each category. **B:** Same shown as a percentage of total reads. White bars: ASA3-p11 reference genome, light grey bars: cells transduced with only one arm, dark grey bars: cells transduced with both arms.

Interestingly, the short wild-type allele, 13 triplets long, also exhibited contractions to 6, 7 and 8 triplets, respectively in clones C6, C3 and C9 (Table 1). This was never observed with the wild-type TALEN. This means that the TALEN_RECODED_ is able to induce a DSB in the short allele, contracting it to an even shorter length.

Finally, in each clone, the occurrence of internal deletions was greatly reduced, confirming the efficacy of our recoding process. All clones contain one to three integration sites for each arm, and at least one of each is intact, except in one ins1tance. In the C11 clone, only the left arm could be detected undeleted (Figure 2B). Note that deletions removed only one or two motifs in the C-terminal part of the protein, and this did not probably affect its capacity to bind to its target and trigger a DSB. This result validated our recoding pipeline for future experiments.

### CTG trinucleotide repeat contractions are not associated with crossovers

In four instances, reads covering the *DMPK* locus exhibited two different CTG repeat lengths for the expanded allele (Table 1). We assumed that these supplemental alleles came from the contraction of the large expanded mutant allele, but we could not formally exclude that they result from an expansion of the wild-type allele. In order to address this question, we used sequence polymorphisms flanking both alleles. The mutant allele carries a G-T transversion 648 bp before the first CTG triplet and an A-C transversion 2286 bp after the last CTG triplet. At least one of the two SNPs could be detected in 84 reads, and both SNPs were identified in 42 reads. None of them could be found in 22 reads, due to low sequence quality (Table 2). Among the reads that were scored as contractions of the mutant allele, none carried a SNP from the WT allele. Conversely, none of the reads assumed to come from the WT allele carried a SNP from the mutant allele. It also shows that the 22 CTG triplets observed in the C9 clone really came from a contraction of the mutant allele and not from a small expansion of the WT allele. Additionally, no recombination of flanking markers was observed in any clone in which both SNPs could be identified. Altogether, this strongly suggests that contractions occur by an intra-allelic mechanism, or by inter-allelic recombination never associated with crossover or long-range conversion of flanking markers.

### Cas9 is less efficient than the TALEN to contract CTG repeats

Meanwhile, it was decided to conduct similar experiments employing the CRISPR-Cas system. Our previous work in *S. cerevisiae* demonstrated that *Streptococcus pyogenes* Cas9 (*Sp*Cas9) was markedly more proficient at inducing a DSB within a CTG repeat tract when compared to *Staphylococcus aureus* Cas9 (*Sa*Cas9) (Poggi *et al* , 2021). Fortunately, both nucleases possess a PAM 3’ of the CTG repeat tract (Figure 5A). However, off-target assessments predict 175 sites (including 40 exonic sites) with 0 mismatches in the human genome for *Sp*Cas9, while *Sa*Cas is associated with only 8 sites (including 6 exonic sites). Given the importance of nuclease specificity in potential gene therapy applications, it was chosen to use *Sa*Cas9, despite its lower efficacy as compared to *Sp*Cas9.

**Figure 5:**
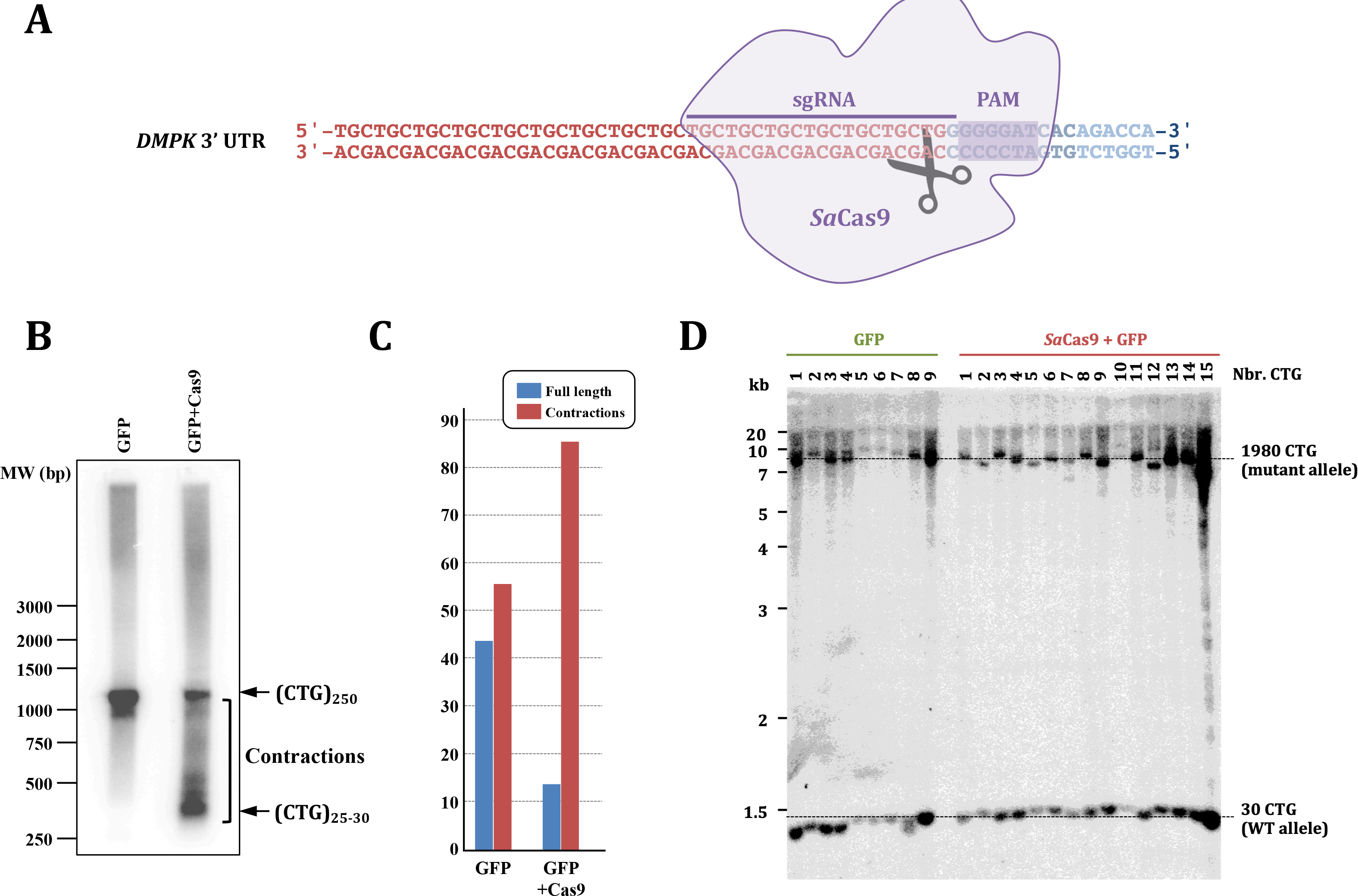
Staphylococcus aureus Cas9 expression in mice and human cells. **A:** Design of the sgRNA and location of the PAM on the 3’ sequence of the *DMPK* gene. **B:** Transduction of GFP alone or *Sa*Cas9, its sgRNA and GFP in heterozygous transgenic mice fibroblasts carrying 250 CTG triplets. The CTG repeat was amplified with primers ST300f and ST300r. **C:** Quantifications of PCR signals from B. **D:** The Southern blot shows independent ASA clones in which the GFP alone or *Sa*Cas9, its sgRNA and GFP were transduced. Although small contractions or expansions could be detected in some clones, no significant differences were found between both experiments.

The gene encoding *Sa*Cas9 along with its associated single-guide RNA (sgRNA) was cloned in lentiviral vector containing the GFP reporter gene, and subsequently transduced into transgenic mice fibroblasts containing a (CTG)_250_ expansion within the human *DMPK* gene. Cells transduced solely with the GFP gene exhibited limited contractions, whereas cells transduced with the nuclease and its corresponding sgRNA displayed substantial contractions (Figure 5B). Quantitative analysis of the signal indicated that 85% of the cells transduced with *Sa*Cas9 exhibited partial or nearly complete contraction of the repeat tract (Figure 5C). The PCR product corresponding to the completely contracted allele was gel purified and sequenced. It shows that only 19 CTG triplets were left, which is the maximum length that was observed in budding yeast after TALEN induction (Richard *et al*, 2014).

The same lentiviral vectors were also transduced in human ASA cells. Multiple clones were expanded, and the length of the repeat tract was assessed using Southern blotting, as previously described. Although small expansions and contractions were observed in cells transduced with the GFP alone or in combination with the nuclease, no significant distinction was discerned between the two transduction strategies (Figure 5D). Based on this series of experiments, it was concluded that *Sa*Cas9 demonstrated a much lower efficacy than the TALEN in inducing contractions within an expanded CTG trinucleotide repeat in mouse cells and proved entirely ineffective in human cells.

## Materials & Methods

### Plasmids

The wild-type TALEN used here is the same as the one used in *S. cerevisiae*, since the CTG trinucleotide repeat integrated in the yeast genome comes from a DM1 patient (Richard *et al*, 2014). First, the right TALEN arm from pCMha188KN16715 (Mosbach *et al* , 2018b) was cloned into pCDNA3.1 (-) (Sigma Aldrich), along with the enhanced GFP gene, to give rise to the pRight plasmid. pMOV1 was built directly in yeast, by homologous recombination between pRight, the left TALEN arm extracted from pCMha182KN9996 (Mosbach *et al* , 2018b) and an *URA3*-CEN-ARS piece of DNA from pRS416 (Sikorski & Hieter, 1989). Both TALEN arms are under the control of the CMV promoter.

For lentiviral expression, the left TALEN arm was cloned into pHR’CMVGFP_hRIF1(1924- 2446)IREShygro3 (Addgene #23138) to give plasmid pTRI204. The right arm was cloned into pLVbcNEOPGK-IRES-Neo to give plasmid pTRI205. For Cas9 lentiviral expression, we used a *Sa*Cas9 and GFP-containing backbone (Addgene #118836, plasmid pTRI211) into which was cloned the *Sa*Cas9-CTG sgRNA to give plasmid pTRI 212.

The two recoded TALEN arms were ordered as synthetic pieces of DNA delivered into the pUC57-Mini vector (GenScript). The left recoded arm was cloned in pHR’CMVGFP_hRIF1(1924-2446)IREShygro3 (Addgene #23138) at BamHI and XhoI to give plasmid pTRI213. The right recoded arm was cloned in pLVbcNEOPGK-IRES-Neo at AgeI and SalI to give plasmid pTRI214. All plasmid maps were created with SnapGene and are shown in Supplemental Figure S3.

### Mice fibroblasts

Fibroblasts were isolated from transgenic mice lungs, carrying a hemizygous (CTG)_600_ expansion from a human DM1 patient (Gourdon *et al*, 1997) (cell line 6423). Transfection was performed using the Viafect commercial reagent (Promega), following manufacturer conditions. Cells were transfected with the pRight plasmid, containing only the right TALEN arm, or with the pMOV1 plasmid, containing both TALEN arms. GFP-positive cells were sorted on the Institut Pasteur cytometry platform. Cells were collected 48 hours after transfection, whole genomic DNA was extracted and analyzed by PCR, using primers ST300f (GAACTGTCTTCGACTCCGGG) and ST300r (GCACTTTGCGAACCAACGAT) as previously described (Tomé *et al*, 2009).

### Human ASA fibroblasts

Immortalized ASA cells (Arandel *et al* , 2017) were grown at 37°C in a 5% CO_2_ humidified atmosphere, in DMEM (Gibco #61965) supplemented with 15% fetal bovine serum (FBS). Cells were passaged by briefly washing with PBS before using trypsin for 5 min at 37°C to dissociate cells, before being split in two flasks. For long-time conservation, cells were frozen in 90% FBS +10% DMSO on dry ice, before being stored in liquid nitrogen. For lentiviral transductions cells were seeded one day before at 55,000 cells/ well in 12-well plates. In the morning, media was changed for 500 µL of DMEM +5% FBS +4 μg/mL of polybrene (Merck TR-1003-G). Lentiviruses were diluted to the desired MOI in 500 µL of DMEM +5% FBS and slowly added to each well. Cells were incubated at 37°C in a 5% CO_2_ humidified atmosphere overnight. The following morning, 1 mL of fresh DMEM +25%FBS was added. The following day, the medium was changed to 500 µL of DMEM +15% FBS, antibiotics selection pressure was added and maintained for 1 week.

### Ring cloning

Cells were diluted to reach between 10 to 100 cells per plate. Media was changed every week until small colonies of cells were detectable under the microscope (usually around 1 month). Media was removed, plastic rings were glued around each colony using silicone. After washing with PBS, trypsin was added inside the ring to dissociate the cells. Colonies were seeded into 96-well plates. Cells were expanded reaching the desired number of cells, by passaging into 24-well plates, 6-well plates, T25 and T75 flask, for an overall culture time of 2-3 months.

### DNA extraction and Southern blotting

Around 1x10^6^ cells were harvested from one T75 flask, washed in PBS and resuspended in 300 µL lysis buffer (100 mM Tris-Hcl pH8.0, 5 mM EDTA pH8.0, 0.2% SDS and 200 mM NaCl) supplemented with 600 µg of proteinase K, and incubated overnight in a water bath at 55°C. In the morning, 300 µl of phenol-CIAA were added to a Phase Lock Gel (Light) microtube (QuantaBio #2302820), following manufacturer instructions. The cell lysate is collected, added to the phenol-CIAA and manually mixed several times (no vortexing). After centrifugating at 13,000 rpm, the aqueous phase was collected, and DNA was extracted using Chloroform-IAA in a new Phase Lock Gel microtube. Collected DNA was subsequently precipitated by adding 1/10 volume of 3M NaCl and 3 volumes of absolute ethanol. Centrifugation at 13,000 rpm was carried out at 4°C for 1 hour. DNA pellet was washed in 70% ethanol and resuspended in TE.

For Southern blotting, 10 µg of DNA were digested with BamHI (60U, NEB) overnight. DNA was loaded in a 0.9% agarose +BET gel, in 1X TBE. Samples were run at 1V/cm, in 1X TBE at 4°C, overnight. A picture of the gel after migration was taken under a BioRad imaging system. The gel was quickly rinsed in distilled water and treated for one hour in 1M NaOH, followed by 2 hours in Tris 1M pH8.5, NaCl 3M. The gel was manually transferred onto a positively charged nylon Hybond-XL membrane (Cytiva #RPN203S), overnight in 6X SSC. The membrane was crosslinked under a Stratalinker at 0.120 J/cm^2^ (Uvitec). Prehybridization was carried tout into Sigma PerfectHybridPlus solution at 68°C for 30 minutes. To prepare the probe, B1.4 plasmid containing a cloned 3’UTR of DMPK gene was digested with BamHI and the 1390 bp band was gel purified. The probe was labeled with alpha-32P CTP using the High Prime DNA labeling kit (Roche, #11585584001). The labelled probe was purified on a G50 sephadex column (Cytiva, #28903408) and denatured for 5 minutes at 95°C. Hybridization was carried out at 68°C overnight. Two 20 min washes were carried out in high stringency buffer (0,5% SSC + 0,1% SDS), followed by one wash in ultra high stringency buffer (0,1% SSC + 0,1% SDS). The membrane was wrapped in plastic and exposed to a phosphorimager screen for 1-2 days (or more if signal was weak).

### Bioinformatics

Off-target predictions for *Sp*Cas9 and *Sa*Cas9 were evaluated using the online CRISPOR tool (Haeussler *et al*, 2016), on the human GRCh38 reference assembly. The OSTINATO pipeline is available on request to S. D.-D. or G.-F. R. and will be extensively described elsewhere. Shortly, the algorithm first detects low complexity zones in HiFi reads and filters out other sequences. It subsequently counts the number of CTG (or CAG) triplets in the six frames within these reads and consider them to be part of a repeat if their number reaches a certain threshold set by the user. This approach dramatically reduces the number of false positives (tandem repeats inadequately identified as CTG/CAG repeats) due to low quality sequences.

### Statistics

Expansions and contractions observed by Southern blot with the TALEN or the TALEN_RECODED_ were compared using a Fisher exact test (Figure 3). Length distributions observed by PacBio sequencing at the *DMPK* locus following TALEN or TALEN_RECODED_ expression were compared using the Student t-test (Figure 4). Confidence intervals for imperfect and perfect repeat length were calculated from the mean of the length distribution (m) and its standard deviation (σ). Repeat lengths outside of m ± 2σ (95% confidence interval) were considered to be statistically different from the mean of the distribution (Table 1). All Statistics were performed using the R package (Millot, 2011).

### PacBio sequencing and analysis

Total genomic DNA was prepared as for Southern blotting, except that after Proteinase K treatment, the remaining pellet debris were not resuspended by pipetting but by gentle taping on the side of the tube, in order to avoid breaking high molecular weight DNA. Subsequent pipetings during phenol-CIAA treatments were performed with blue cones. PacBio libraries were prepared on the Biomics platform (Institut Pasteur), as follows. Genomic DNAs of clones in which the WT TALEN was transduced (ASA30, ASA51, ASA2, ASA53, ASA54, ASA27, ASA111E, ASA22C, ASA28 and ASA03-44, Supplemental Table S2) were sheared in g-tubes (Covaris) to an average of 50 kb length. Libraries were prepared with the SMRTbell Express Template Prep Kit 2.0 (Pacific Biosciences). Sequences were performed on the Sequel PacBio machine at the Pasteur Biomics platform using the Sequel Binding Kit 3.0 and the Sequel DNA Internal Control 3.0 (Pacific Biosciences). Genomic DNAs of the reference genome (ASA3-p11) and of clones in which the TALEN_RECODED_ was transduced (C3, C6, C9, C11, C12, C15, C16, C3_Left and C5_Left, Supplemental Table S2) were sheared in g-tubes to an average of 15 kb length. Libraries were prepared with a SMRTbell Express Template Prep Kit 2.0 (Pacific Biosciences). Sequences were performed on a Sequel II machine, at the Institut Curie ICGex NGS platform. Loading concentration of each sample was 6 pM and run time was 20 hours for both sets of clones.

Quality control and primary assembly of the ASA3-p11 reference genome was performed with QUAST-LG, a versatile assembler that can work with short or long reads (Mikheenko *et al* , 2018).

## Discussion

Myotonic dystrophy type 1 is triggered by the expansion of a CTG trinucleotide repeat within the 3’ UTR of the *DMPK* gene, located on the human chromosome 19. In the present study, we have demonstrated the remarkable efficacy of a TALEN in contracting the expanded repeat in transgenic mice cells and human DM1 cells. Previous therapeutic strategies for treating DM1 have primarily focused on inhibiting *DMPK* expression (Pascual-Gilabert *et al* , 2021), or targeting the CUGexp-mRNA to prevent its interaction with the MBLN1 protein (reviewed in Izzo *et al* , 2022). However, thus far, none of these approaches has advanced beyond clinical trial stages.

At the DNA level, various attempts have been made to completely remove the CTG repeat tract by inducing double-strand breaks upstream and downstream using the CRISPR-Cas9 system. Enhanced *Streptococcus pyogenes* Cas9 (e*Sp*Cas9) was transfected into immortalized myoblasts harboring 300-1,000 CTG triplets within the *DMPK* locus. Among the 85 clones examined, only 12 exhibited a pattern consistent with the complete deletion of one or two alleles. Seven out of these 12 clones exhibited a marked reduction in nuclear foci (Provenzano *et al* , 2017). PCR amplifications were conducted on seven putative off-target sites to confirm the absence of other genomic modifications.

In another study, wild-type *Sp*Cas9 was expressed in iPSC derived from DM1 patients with the same objective (Dastidar *et al* , 2018). Among 25 independent clones analyzed, 19 demonstrated deletions of the CTG repeat tract. However, nine of these exhibited deletions larger than the intended target locus, resembling observations made in *S. cerevisiae* cells using the same nuclease and a similar approach (Mosbach *et al* , 2020). *Sp*Cas9 was also expressed in DM1 patient myoblasts, which carried 2,600 CTG triplets on the expanded mutant allele and 13 triplets on the wild-type allele. Among the 103 myoblasts clones examined, 35% exhibited deletions of the repeat tract on both alleles, while 51% showed deletions solely on the mutant allele (Agtmaal *et al* , 2017). Notably, neither of these last two reports conducted an analysis of putative off-target sites.

Lo Scrudato and colleagues utilized *Staphylococcus aureus* Cas9 (*Sa*Cas9), a smaller nuclease compared to *Sp*Cas9 and more suitable for co-packaging with the sgRNA within the same viral particle (Lo Scrudato *et al* , 2019). They expressed *Sa*Cas9 in DM1 myoblasts carrying 2,600 CTG triplets on the mutant allele. Fifty independent clones expressing the nuclease were isolated. Among these clones, five did not exhibit significant nuclear foci. Subsequently, the authors expressed the same nuclease in muscle fibers of transgenic mice harboring a large CTG expansion (1,200 CTG triplets) at the *DMPK* locus integrated in their genome (Gomes-Pereira *et al* , 2007). PCR analysis conducted on muscle fibers suggested that approximately 10% of the nuclei contained a deletion of the repeat tract. Unfortunately, no analysis of putative off-target sites was conducted.

Alternative strategies directed at modifying DNA include the insertion of a polyA signal upstream of the CTG expansion through TALEN-driven homologous recombination (Xia *et al*, 2015) and the utilization of a Zinc-Finger Nuclease (ZFN) or *Sp*Cas9 to contract CTG repeats in a HEK293 cell model containing a GFP-reporter gene that becomes active upon contraction of the CTG repeat tract (Cinesi *et al* , 2016). This latter approach closely resembles what we proposed a decade ago using a TALEN (Richard *et al* , 2014). In the HEK293 experimental system, the authors observed expansions and contractions upon the introduction of a ZFN or *Sp*Cas9 targeting the CTG trinucleotide repeat. Notably, these events were more frequent when employing ZFN as compared to *Sp*Cas9. Intriguingly, when expressing a *Sp*Cas9 D10A mutant in which one of the nuclease activities was disabled, expansions occurred with a reduced frequency, and a four-fold increase in contractions was reported.

### A TALEN is very efficient at contracting an expanded CTG repeat at the human DMPK locus

In the present study, we applied the previously successful approach used in budding yeast to contract an expanded CTG repeat. The initial wild-type TALEN (Richard *et al*, 2014) exhibited high efficiency in murine fibroblasts carrying 600 CTG triplets. However, when the same TALEN was transduced in human cells carrying 1800 triplets, we observed less frequent contractions, of limited magnitude (Figure 1). Moreover, internal deletions occurring via homologous recombination between TALE repeats were identified in each TALEN arm. The precise timing of these deletions remains unclear, whether they happened during reverse transcription of the lentiviral RNA genome or after integration in the ASA genome. Nevertheless, it is plausible to hypothesize that the TALEN had been actively expressed for some duration before becoming inactivated due to deletions. This hypothesis is supported by the observation of contractions occurring at the *DMPK* locus and other CTG repeats which were les frequent when the same cells were transduced with the right arm alone.

By recoding the TALEN using the degeneracy of the genetic code, we were able to introduce two significant modifications. First, the recoding reduced the sequence similarity between motifs from 94%-100% to 65%-78%. This resulted in the retention of the TALEN as full-length arms in each clone analyzed, except C11 (Figure 2). It is worth noting that recoded TALENs developed by the Church lab have previously demonstrated comparable efficacy to wild-type TALENs in inducing homologous recombination at the AAVS1 locus in hiPSCs (Yang *et al*, 2013). These recoded TALENs could also be efficiently packaged into lentiviral particles and transduced into HEK293T cells without compromising the sequence integrity of the TALE repeats. In our case, the TALEN_RECODED_ was engineered to ensure that no more than 20 nucleotides of identity remained between two TALE motifs. Furthermore, rare codons were eliminated to optimize its expression in human cells. This customized TALEN_RECODED_ outperformed the original TALEN in inducing CTG repeat contraction at the DMPK locus (Table 1 and Figure 4). In addition, it reduced the incidence of homologous recombination between TALE repeats, thus ensuring the preservation of full-length arms. Of particular interest, the TALEN_RECODED_ substantially decreased the occurrence of off-target contractions in other long CTG repeats. This observation highlights that enhancing on-target efficacy does not necessarily lead to an increase in undesired off-target mutations. Mutations in other CTG trinucleotide repeats fall in one intergenic region and one intron, in the *MYO9A* gene (Table 1). It is therefore reasonable to assume that their impact on gene function will be negligible. The clonal cell lines containing these contractions did not show any obvious growth defects, as compared to non-transduced cells.

Notably, *Sa*Cas9 displayed lower efficacy compared to the TALEN in triggering CTG repeat contractions in transgenic mice cells and was entirely ineffective in ASA cells (Figure 5). This lack of efficacy in inducing DSBs within CTG repeats mirrors observations made in *S. cerevisiae* when using the same repeats and the same nuclease. This highlights the extreme value of the yeast model in assessing the efficacy of forthcoming DNA endonucleases (Poggi *et al*, 2021).

### Molecular mechanisms underlying trinucleotide repeat contractions

The examination of SNPs bordering the CTG repeat tract revealed that contractions were not associated to crossovers or long-range gene conversion events, such as those stemming from Break-Induced Replication (BIR). Consequently, this observation strongly indicates that contractions likely occur through an intra-allelic mechanism, such as Single-Strand Annealing (SSA). In budding yeast, our investigations demonstrated that TALEN-induced CTG repeat contractions implicated the Mre11-Rad50 complex, along with the Sae2 and Rad52 proteins. Notably, the deletion of the *RAD51* gene (RecA homologue), the BIR-driving *POL32* gene, or *DNL4* , the ligase central to end-joining, had no impact on the DSB-repair of the CTG repeat tract (Mosbach *et al* , 2018b). Consequently, we concluded that contractions occured via SSA, an intra-allelic mechanism necessitating solely the Mre11-Rad50 complex and Rad52. This mechanism is compatible with our present observations, leading us to propose that CTG repeat contractions via DSB repair also occur through SSA in human cells.

Although CTG repeats generally exhibit expansion across successive generations, sporadic instances of contractions have been documented. In a family where the father harbored a 600 CTG expansion at the DMPK locus, one of his daughters inherited a nearly complete contraction, reducing the count to 13 triplets, while remaining asymptomatic. Examination of flanking genetic markers led the authors to posit that this contraction resulted from a discontinuous gene conversion event involving both chromosomes (O’Hoy *et al*, 1993). In our study, recombination of flanking SNPs was never observed; however, we cannot entirely rule out the possibility that contractions occurred via Synthesis-Dependent Strand Annealing (SDSA), a gene conversion mechanism not previously associated with crossovers but demonstrated to induce tandem repeat instability during DSB repair in budding yeast (Pâques *et al*, 1998, 2001; Richard *et al*, 2003, 2000; Richard & Pâques, 2000).

It is noteworthy that the left arm alone triggers repeat contractions in ASA cells, a surprising finding that raises several intriguing possibilities. One hypothesis is that it may self-dimerize on the CTG repeat, potentially causing a DSB, as previously proposed by others (Liu *et al* , 2010). Alternatively, it is possible that the left arm induces a single-strand break (nick) that proves sufficient to trigger repeat contractions. Notably, one single FokI domain has been demonstrated to induce nicks on a supercoiled plasmid (Bitinaite *et al* , 1998). This latter hypothesis is compatible with findings from the Dion lab which showed that the *Sp*Cas9 nickase induces CTG repeat contractions in HEK293T cells (Cinesi *et al* , 2016). Another possibility is that the binding of the left arm to the CTG repeat tract might trigger contractions by interfering with replication during growth of cell cultures. This effect would be lower with the right arm alone, as it binds to the *DMPK* locus in only one copy, whereas the left arm may bind multiple times along the expanded CTG triplet repeat.

Future directions for this research encompass enhancing the TALEN specificity. This can be achieved by using different FokI domains (Miller *et al*, 2007), and examining alternative RVD for the TALE repeats. Furthermore, it is imperative to delve into the precise mechanism by which the left arm, when used alone, triggers CTG repeat contractions. Understanding this mechanism, holds the potential to provide a viable alternative to DSBs in the long-term vision of establishing an efficient gene therapy approach. Such an approach could be applied not only for DM1 but also to address all other microsatellite expansion disorders.

## Acknowledgements

1. L. B. was supported by a PasteurInnov grant to G.-F. R. (project NUCMYO). O. F. was supported by an AFM-Telethon grant to G.-F. R. (project 21431). L. P. was supported by a PhD Cifre fellowship from SANOFI. V. M. was supported by a PhD fellowship from the french Ministère de l’Enseignement Supérieur et de la Recherche. PacBio sequencing was very generously funded by the Groupama foundation. Biomics Platform, C2RT, Institut Pasteur and ICGex–NGS Platform, Curie, Paris, France are supported by France Génomique (ANR-10-INBS-09) and IBISA. This work was supported by the AFM-Telethon, the Institut Pasteur, the Association Institut de Myologie, and the CNRS.

We acknowledge the help of the HPC Core Facility of Institut Pasteur for this work. We also wish to thank Pierre-Henri Commere for efficient cell sorting on the cytometry platform of the Institut Pasteur and the vectorology platform of the Institut du Cerveau et de la Moelle (CHU La Pitié-Salpétrière, Paris) for lentivirus productions.

## Author contributions

B.D., G.G., D.F. and G.-F.R. conceived the experiments; L.B., O.F., L.P., V.M., S.T., D.V. and A.K. carried out experiments with TALEN and Cas9; L.M., S.L., T.C. and M.M. carried out PacBio sequencing; S.D.D. and G.-F.R. analyzed the data; G.-F.R., G.G. and D.F. wrote the article.

## Sequence deposit

The ASA3-p11 DM1 reference genome sequence was deposited at the NCBI (XXX-000)

---

Supplemental Figure S1

Correlation between CTG repeat length measured on Southern blot (X axis) and determined by PacBio sequencing (Y axis). Lengths are in nucleotides.

Supplemental Figure S2

**Top** : Sequence comparison of TALE repeats between the original wild-type TALEN and the TALEN_RECODED_. Red rectangles show sequence stretches at least 3 nucleotide long that were conserved between the original and the recoded motif. The two large black rectangles underline the six nucleotides encoding the RVD. **Bottom** : Multiple alignments of the 15 TALE motifs of each arm, for the TALEN and the TALEN_RECODED_. Alignments were performed using Clustal Omega, minimum and maximum identity percentages between two motifs were calculated by MView. Both programs were run on the NCBI-EBI multiple alignment portal (https://www.ebi.ac.uk/Tools/msa/).

Supplemental Figure S3

Plasmids used in this study, all available on request to G.-F.R.

Supplemental Table S1

Sequencing data of the DM1 reference genome (ASA3-p11)

Supplemental Table S2

Summary of PacBio sequences performed on Sequel or Sequel II machines. For each clone, TALEN arms, and the number of SMRTCells are shown.

Supplemental Table S3

For each clone, the total amount of sequence recovered after PacBio sequencing is shown, along with the genome coverage and the percentage of genome covered.

## Notes

### Competing Interest Statement

The authors have declared no competing interest.

### Summary of Updates

Sonia Lameiras added as an author New Figure 2 New PacBio sequences were analyzed and added to Table 1 and Figure 4 Material and Methods completed with statistics and computer analysis methods

