## Supplementary material for "TALEN-induced contraction of CTG trinucleotide repeats in myotonic dystrophy type 1 cells": Table 1

**Table 1: Median length of perfect and imperfect CTG repeats as determined by PacBio sequencing**

Grey numbers identify length differences outside the 95% confidence interval, as compared to the reference genome

Lengths are in triplets, except for TRF for which they are in nucleotides

**WILD-TYPE TALEN ($TALEN_{WT}$)**

| **Repeat consensus** | Imperfect CTG | | Imperfect CTG | | Imperfect CTG | | Imperfect CTG | | Imperfect CTG | | Imperfect CTG | | Imperfect CTG | | Imperfect CTG | | Perfect (CTG)58 | | DMPK CTG expansion | |
|---|---|---|---|---|---|---|---|---|---|---|---|---|---|---|---|---|---|---|---|---|
| **Chromosome** | **2** | | **3** | | **4** | | **6** | | **8** | | **10** | | **15** | | **15** | | **17** | | **19** | |
| Repeat ID# | 2p13_1 | | 3p25 | | 4q35 | | 6q25 | | 8q24 | | 10q24 | | 15q11 | | 15q23_2 | | 17q21_2 | | 19q13 | |
| Gene | *SNRNP27* | | *TPRXL* | | - | | *SLC22A22* | | - | | - | | - | | *MYO9A* | | *CA10* | | *DMPK* | |
| Location | Intron | | Intron | | Intergenic | | Intron | | Intergenic | | Intergenic | | Intergenic | | Intron | | Intron | | 3' UTR | |
| ASA3_p11 (reference) | 86 | | 128 | | 111 | 79 | 48 | 47 | 64 | | 56 | | 278 | 256 | 89 | | 58 | | 1806 | 13 |
| ASA03-44 | 86 | | 128 | | 111 | 79 | 48 | 47 | 63 | | 56 | | 279 | 256 | 89 | | 61 | | 1879 | 14* |
| ASA2 | ND | | ND | | 108* | | 38 | | 62* | | ND | | | 206* | 88 | | 60 | | 1235* | 13 |
| ASA22C | 82 | | 125 | | | 72* | 27* | | 58 | | 46 | | | 252* | 80* | | 52* | | 1745* | 13* |
| ASA27 | 86 | | 128 | | 111 | 79 | 48 | 47 | 64 | | 56 | | 278 | 257/227 | 89 | | 59 | | 1272/207* | 13 |
| ASA28 | 78 | | 127* | 112* | 105* | | ND | | ND | | 48 | | ND | | 89* | | ND | | ND | 13* |
| ASA30 | 81* | | ND | | | 66 | ND | | ND | | ND | | | 251 | 83 | 71 | 60* | 47* | ND | 11* |
| ASA51 | 83 | | 126 | | 110* | | ND | | 63* | | 55 | | ND | | ND | | 59* | | 2278* | ND |
| ASA53 | 86 | | 128 | | 111 | 79 | 48 | 47 | 64 | | 56 | | 278 | 256 | 89 | | 61 | | ND | 13 |
| ASA54 | 86 | | 128 | | 111 | 80 | 48 | | 64 | | 56 | | 278 | 256 | 89 | | 61 | 56 | 1471 | 12 |
| ASA111E | 79 | | ND | | ND | ND | 47 | 39 | 58* | | 45 | | 276 | 256 | ND | | 55 | | ND | 13 |
| Found by TRF | 148 | | 200 | | 116 | 158 | 84 | 109 | ND | | 147 | | ND | | 157 | | 64 | | 1766 | ND |
| Contractions (14) | | | | 1 | 1 | 1 | 2 | 1 | 2 | | 3 | | | 2 | | | | | 1 | |
| Expansions (1) | | | | | | | | | | | | | | | | | | | 1 | |

**RECODED TALEN ($TALEN_{recoded}$)**

| **Repeat consensus** | Imperfect CTG | | Imperfect CTG | | Imperfect CTG | | Imperfect CTG | | Imperfect CTG | | Imperfect CTG | | Imperfect CTG | | Imperfect CTG | | Perfect (CTG)61 | | DMPK CTG expansion | |
|---|---|---|---|---|---|---|---|---|---|---|---|---|---|---|---|---|---|---|---|---|
| **Chromosome** | **2** | | **3** | | **4** | | **6** | | **8** | | **10** | | **15** | | **15** | | **17** | | **19** | |
| Repeat ID# | 2p13_1 | | 3p25 | | 4q35 | | 6q25 | | 8q24 | | 10q24 | | 15q11 | | 15q23_2 | | 17q21_2 | | 19q13 | |
| Gene | *SNRNP27* | | *TPRXL* | | - | | *SLC22A22* | | - | | - | | - | | *MYO9A* | | *CA10* | | *DMPK* | |
| Location | Intron | | Intron | | Intergenic | | Intron | | Intergenic | | Intergenic | | Intergenic | | Intron | | Intron | | 3' UTR | |
| ASA3_p11 (reference) | 86 | | 128 | | 111 | 79 | 48 | 47 | 64 | | 56 | | 278 | 256 | 89 | | 58 | | 1806 | 13 |
| C3_Left (left arm) | 86 | | 128 | | 111 | 79 | 48 | 47 | 64 | | 56 | | 278 | 255 | 88 | | 58 | | 41 | 13 |
| C5_Left (left arm) | 86 | | 128 | | 113 | 74 | 48 | 46 | 63 | | 56 | | ND | ND | 88 | 53 | 61 | | 395/190 | 12 |
| C3 | 86 | | 128 | | 111 | 79 | 48 | 47 | 63 | | 56 | | 253 | 225 | 89 | | 57 | | ND | 7 |
| C6 | 86 | | 128 | | 111 | 79 | 48 | 47 | 63 | | 56 | | 278 | 236 | 89 | | 60 | | 489/264* | 6 |
| C9 | ND | | 128 | | 111 | 79 | 48 | 45* | 63 | | 56 | | | 229* | 87 | | 62 | | 22* | 8 |
| C11 | 86 | | 128 | | 111 | 79 | 48 | 46 | 64 | | 56 | | | 255 | 89 | | 58 | | 432 | ND |
| C12 | 86 | | 128 | | 111 | 79 | 48 | 47 | 63 | | 56 | | 274 | 257 | 89 | | 57 | | 517/203* | 11 |
| C15 | 86 | | 128 | | 111 | 79 | 48 | 47 | 63 | | 56 | | 258 | 240 | 89 | | 58 | | 126* | 10 |
| C16 | 86 | | 128 | | 111* | 79* | 48 | 46 | 64 | | 56 | | ND | ND | 89* | | 61 | | ND | 8* |
| Contractions (20) | | | | | | | | | | | | | 2 | 4 | | 1 | | | 10 | 3 |
| Expansions (0) | | | | | | | | | | | | | | | | | | | | |

* repeat entirely covered by only one HiFi read

ND: Not Detected
