## Supplementary material for "TALEN-induced contraction of CTG trinucleotide repeats in myotonic dystrophy type 1 cells": Table 2

**Table 2: Complete list of HiFi reads covering the *DMPK* locus, showing upstream and downstream SNPs flanking the CTG repeat tract**

| Experiments with the $TALEN_{WT}$ | | | | | |
|---|---|---|---|---|---|
| Clone | HiFi read ID | Mutant allele | WT allele | upstream SNP | downstream SNP |
| ASA3p11 | 45157001 | 1930 | | T (Mutant) | - |
| ASA3p11 | 167970116 | 1806 | | T (Mutant) | - |
| ASA3p11 | 174590397 | 1735 | | T (Mutant) | C (Mutant) |
| ASA3p11 | 69732891 | | 14 | G (WT) | A (WT) |
| ASA3p11 | 107610886 | | 13 | G (WT) | A (WT) |
| ASA3p11 | 9241536 | | 13 | G (WT) | A (WT) |
| ASA3p11 | 14288584 | | 13 | - | A (WT) |
| ASA3p11 | 36309499 | | 13 | - | - |
| ASA3p11 | 51839863 | | 13 | G (WT) | A (WT) |
| ASA3p11 | 98044693 | | 13 | G (WT) | A (WT) |
| ASA3p11 | 19530859 | | 12 | G (WT) | A (WT) |
| ASA03-44 | 77070746 | 1913 | | T (Mutant) | C (Mutant) |
| ASA03-44 | 135200843 | 1845 | | T (Mutant) | C (Mutant) |
| ASA03-44 | 145425490 | | 14 | G (WT) | A (WT) |
| ASA2 | 12452809 | 1235 | | T (Mutant) | C (Mutant) |
| ASA2 | 16122728 | | 14 | G (WT) | A (WT) |
| ASA2 | 18940258 | | 12 | G (WT) | - |
| ASA22C | 58000002 | 1745 | | - | - |
| ASA22C | 53805120 | | 13 | - | - |
| ASA27 | 123076975 | 1571 | | T (Mutant) | C (Mutant) |
| ASA27 | 17760336 | 1454 | | - | - |
| ASA27 | 124257412 | 1272 | | - | C (Mutant) |
| ASA27 | 1050798 | 1268 | | T (Mutant) | C (Mutant) |
| ASA27 | 167839782 | 1249 | | T (Mutant) | - |
| ASA27 | 139919677 | 207 | | T (Mutant) | - |
| ASA27 | 140511323 | | 14 | G (WT) | A (WT) |
| ASA27 | 5768120 | | 13 | - | - |
| ASA27 | 138217022 | | 13 | G (WT) | - |
| ASA27 | 161481692 | | 13 | - | A (WT) |
| ASA27 | 168165530 | | 13 | - | A (WT) |
| ASA27 | 19269399 | | 13 | G (WT) | A (WT) |
| ASA27 | 49743819 | | 13 | G (WT) | A (WT) |
| ASA27 | 56821774 | | 13 | G (WT) | A (WT) |
| ASA27 | 61475585 | | 13 | G (WT) | A (WT) |
| ASA27 | 123864069 | | 13 | G (WT) | - |
| ASA27 | 84541614 | | 13 | G (WT) | A (WT) |
| ASA27 | 143133120 | | 13 | G (WT) | - |
| ASA27 | 89457225 | | 13 | G (WT) | A (WT) |
| ASA27 | 128649138 | | 12 | G (WT) | - |
| ASA28 | 45220143 | | 13 | G (WT) | - |
| ASA30 | 9830947 | | 11 | - | - |
| ASA51 | 60818321 | 2278 | | T (Mutant) | - |
| ASA53 | 122030747 | | 13 | G (WT) | A (WT) |

| | | | | | |
|---|---|---|---|---|---|
| ASA53 | 113051842 | | 13 | G (WT) | A (WT) |
| ASA53 | 104467222 | | 13 | G (WT) | A (WT) |
| ASA53 | 83954510 | | 13 | G (WT) | A (WT) |
| ASA54 | 27853746 | 1783 | | - | C (Mutant) |
| ASA54 | 52757128 | 1158 | | T (Mutant) | |
| ASA54 | 130286309 | | 13 | G (WT) | - |
| ASA54 | 69862020 | | 11 | - | - |
| ASA111E | 14352993 | | 13 | G (WT) | A (WT) |
| ASA111E | 69862014 | | 13 | G (WT) | - |
| ASA111E | 41091642 | | 12 | - | - |

| Experiments with the $TALEN_{recoded}$ | | | | | |
|---|---|---|---|---|---|
| **Clone** | **HiFi read ID** | **Mutant allele** | **WT allele** | **upstream SNP** | **downstream SNP** |
| C3_gauche | 31720565 | 43 | | T (Mutant) | C (Mutant) |
| C3_gauche | 130416913 | 42 | | T (Mutant) | - |
| C3_gauche | 178915251 | 41 | | T (Mutant) | - |
| C3_gauche | 59507020 | 40 | | - | - |
| C3_gauche | 21627398 | 38 | | - | - |
| C3_gauche | 96993606 | 32 | | T (Mutant) | C (Mutant) |
| C3_gauche | 71828075 | | 13 | G (WT) | A (WT) |
| C3_gauche | 97978570 | | 13 | G (WT) | A (WT) |
| C3_gauche | 68355907 | | 13 | - | - |
| C3_gauche | 32900477 | | 13 | G (WT) | A (WT) |
| C3_gauche | 135006873 | | 13 | G (WT) | - |
| C3_gauche | 74844072 | | 12 | - | - |
| C3_gauche | 158074528 | | 12 | - | - |
| C5_gauche | 128844008 | 421 | | - | C (Mutant) |
| C5_gauche | 67700696 | 395 | | - | C (Mutant) |
| C5_gauche | 61998061 | 364 | | T (Mutant) | C (Mutant) |
| C5_gauche | 151913851 | 190 | | T (Mutant) | C (Mutant) |
| C5_gauche | 126879216 | | 13 | - | A (WT) |
| C5_gauche | 109445691 | | 11 | - | - |
| C3 | 57212975 | | 7 | - | A (WT) |
| C3 | 22022506 | | 7 | - | A (WT) |
| C6 | 7144827 | 572 | | T (Mutant) | - |
| C6 | 66259103 | 540 | | T (Mutant) | - |
| C6 | 67701151 | 489 | | - | C (Mutant) |
| C6 | 146343411 | 459 | | T (Mutant) | - |
| C6 | 125305217 | 432 | | T (Mutant) | - |
| C6 | 85787784 | 264 | | - | - |
| C6 | 93062831 | | 10 | - | - |
| C6 | 11928821 | | 9 | G (WT) | A (WT) |
| C6 | 17369305 | | 6 | - | - |
| C6 | 132909524 | | 6 | G (WT) | A (WT) |
| C6 | 158466540 | | 5 | - | - |
| C9 | 127205668 | 22 | | T (Mutant) | C (Mutant) |
| C9 | 75891089 | | 8 | G (WT) | A (WT) |
| C9 | 123537897 | | 8 | - | A (WT) |

| | | | | | |
|---|---|---|---|---|---|
| C11 | 168363161 | 438 | | - | C (Mutant) |
| C11 | 121112144 | 425 | | T (Mutant) | C (Mutant) |
| C12 | 118883019 | 531 | | T (Mutant) | C (Mutant) |
| C12 | 94896394 | 517 | | - | C (Mutant) |
| C12 | 168888410 | 448 | | T (Mutant) | - |
| C12 | 157026771 | 203 | | T (Mutant) | C (Mutant) |
| C12 | 118228921 | | 13 | - | A (WT) |
| C12 | 25297899 | | 13 | G (WT) | - |
| C12 | 99748544 | | 13 | G (WT) | A (WT) |
| C12 | 113903673 | | 8 | - | - |
| C12 | 18745301 | | 7 | - | - |
| C12 | 126421532 | | 7 | G (WT) | A (WT) |
| C15 | 22612336 | 126 | | - | C (Mutant) |
| C15 | 70976652 | | 11 | - | - |
| C15 | 56690741 | | 10 | - | A (WT) |
| C15 | 166201494 | | 10 | G (WT) | A (WT) |
| C15 | 121506494 | | 6 | - | A (WT) |
| C16 | 68814304 | | 8 | G (WT) | - |
