## Supplementary figures and images for "TALEN-induced contraction of CTG trinucleotide repeats in myotonic dystrophy type 1 cells"

### Supplemental Figure S1

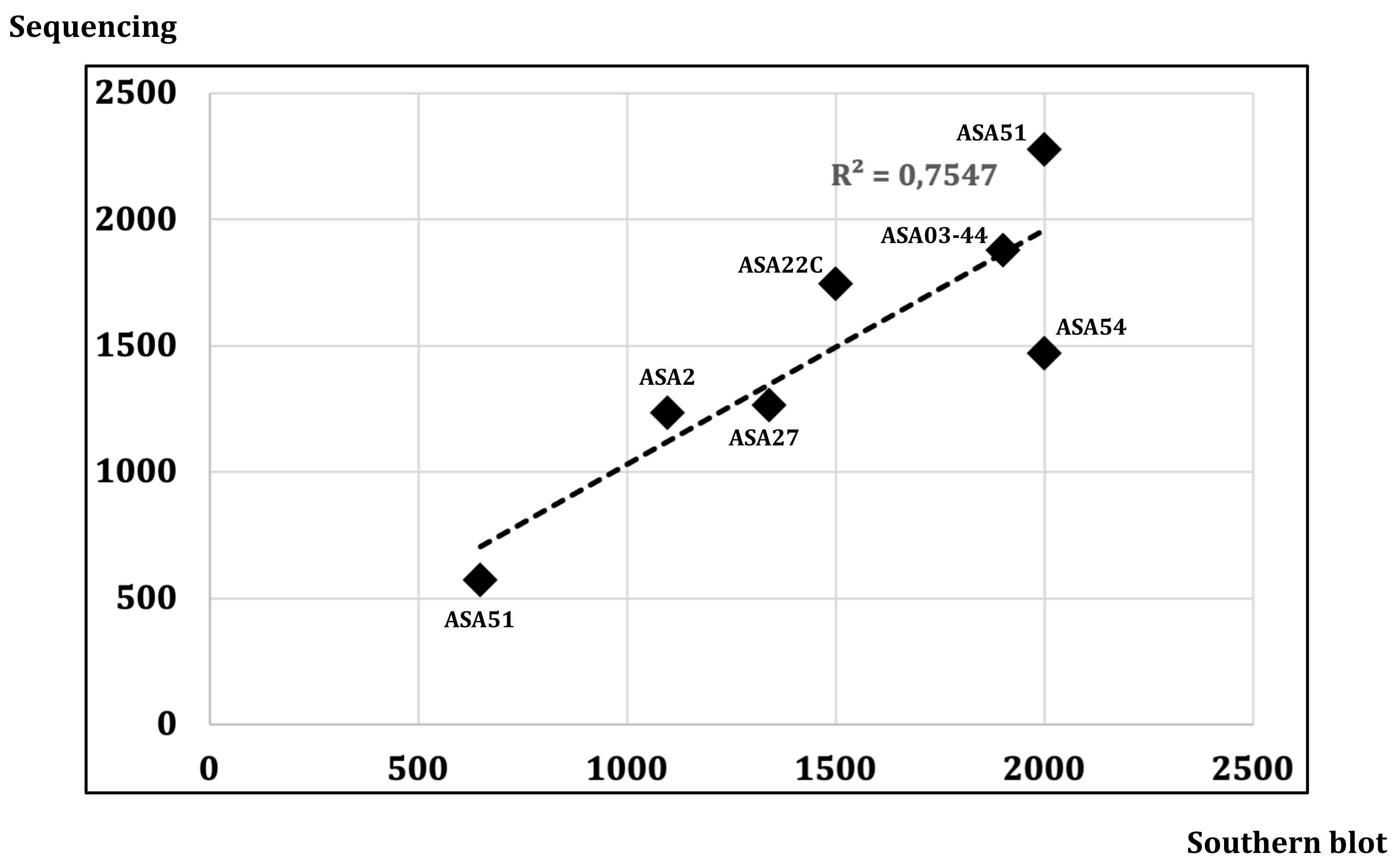

### Supplemental Figure S3

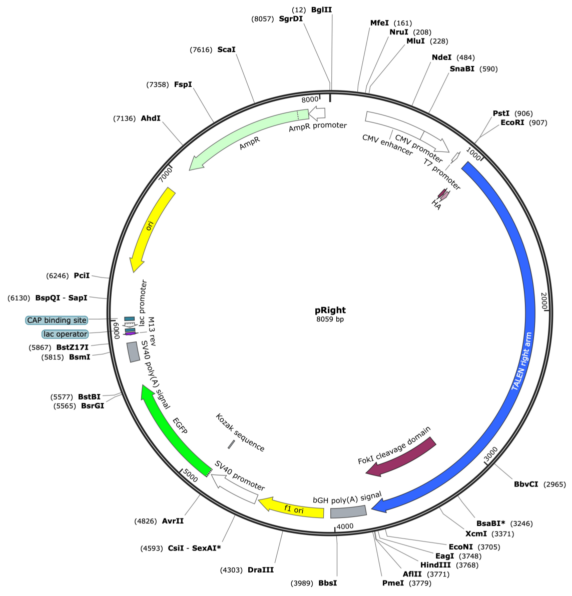

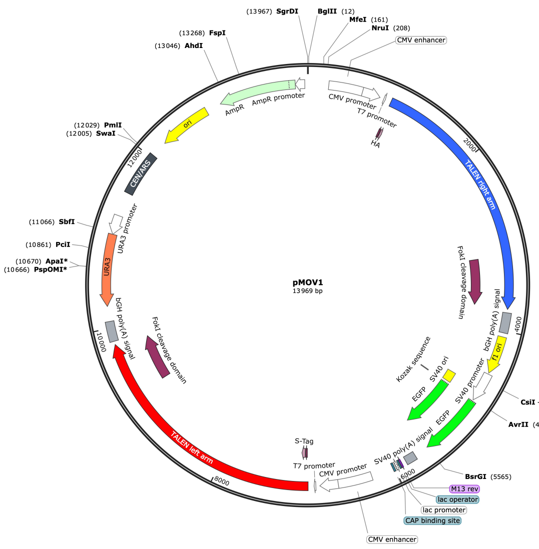

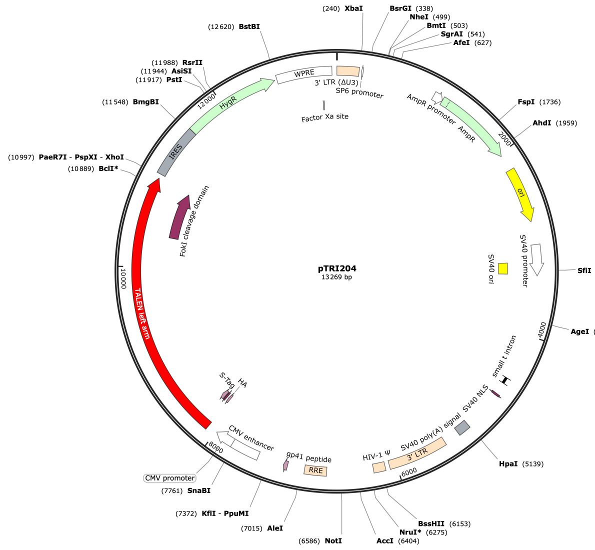

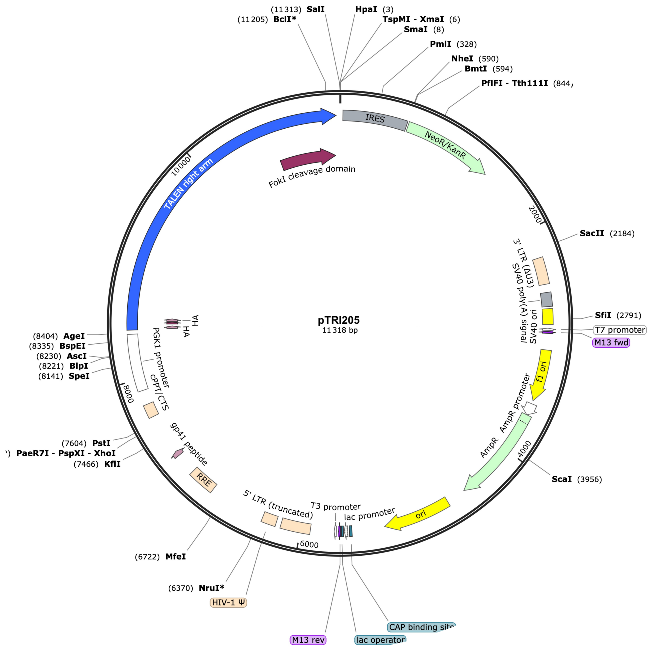

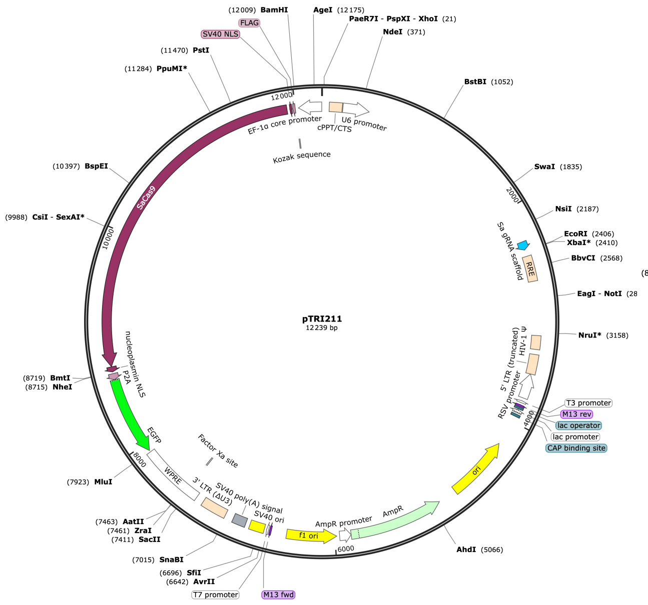

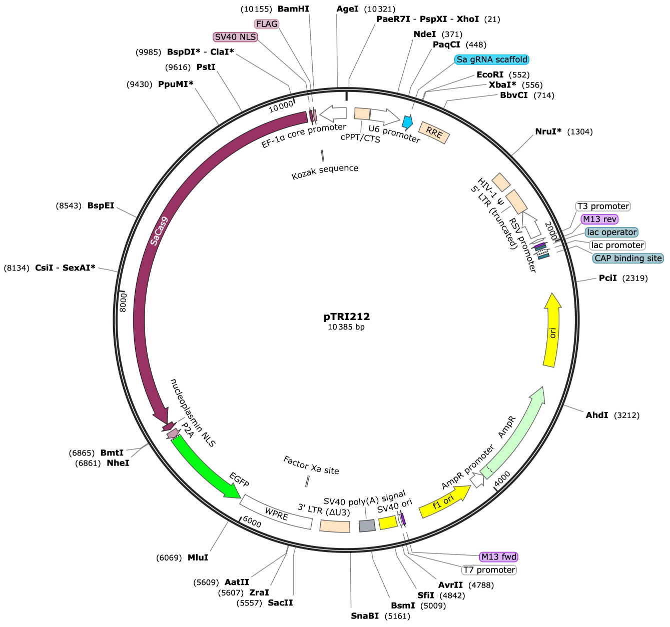

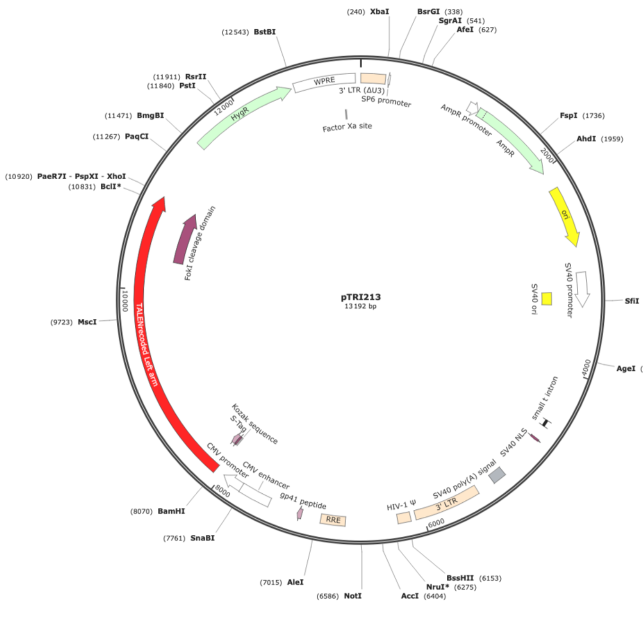

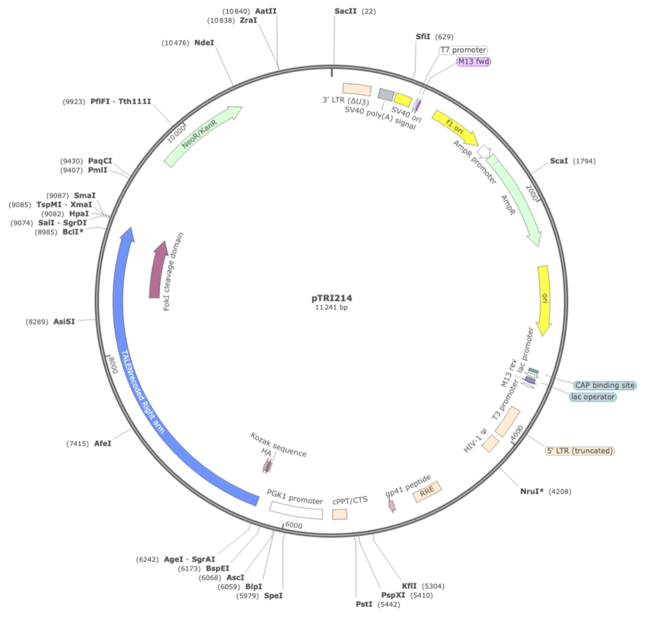
