## Supplemental Figure S2 for "TALEN-induced contraction of CTG trinucleotide repeats in myotonic dystrophy type 1 cells"

Left TALEN arm Binding domain: $G_1C_1T_1G_2C_2T_2G_3C_3T_3G_4C_4T_4G_5C_5T_5$

>$G_1$ (original) TTGACCCCCCAGCAGGTGGTGGCCATCGCCAGCAATAATGGTGGCAAGCAGGCGCTGGAGACGGTCCAGCGGCTGTTGCCGGTGCTGTGCCAGGCCCACGGC
>$G_1$ (recoded) TTGACACCTCAACAAGTTGTTGCTATTGCATCAAATAACGGAGGTAAGCAGGCACTGGAGACCGTTCAGCGGCTGCTCCCTGTGCTTTGTCAAGCTCATGGT
>$C_1$ (original) TTGACCCCGGAGCAGGTGGTGGCCATCGCCAGCCACGATGGCGGCAAGCAGGCGCTGGAGACGGTCCAGCGGCTGTTGCCGGTGCTGTGCCAGGCCCACGGC
>$C_1$ (recoded) CTTACTCCAGAACAAGTCGTCGCAATCGCTTCCCATGACGGGGGCAAACAGGCCCTTGAGACTGTCCAACGCTTGTTGCCAGTTTTGTGCCAGGCACACGGA
>$T_1$ (original) TTGACCCCCCAGCAGGTGGTGGCCATCGCCAGCAATGGCGGTGGCAAGCAGGCGCTGGAGACGGTCCAGCGGCTGTTGCCGGTGCTGTGCCAGGCCCACGGC
>$T_1$ (recoded) CTGACTCCCCAACAGGTGGTAGCTATCGCCTCTAACGGTGGCGGAAAGCAAGCTTTGGAGACAGTCCAAAGGCTCCTGCCCGTTCTCTGCCAGGCCCACGGC
>$G_2$ (original) TTGACCCCCCAGCAGGTGGTGGCCATCGCCAGCAATAATGGTGGCAAGCAGGCGCTGGAGACGGTCCAGCGGCTGTTGCCGGTGCTGTGCCAGGCCCACGGC
>$G_2$ (recoded) CTTACACCACAACAAGTTGTTGCTATTGCTAGTAACAATGGCGGAAAACAAGCCCTGGAGACTGTGCAGCGGCTCCTTCCTGTCCTTTGTCAGGCTCATGGG
>$C_2$ (original) TTGACCCCGGAGCAGGTGGTGGCCATCGCCAGCCACGATGGCGGCAAGCAGGCGCTGGAGACGGTCCAGCGGCTGTTGCCGGTGCTGTGCCAGGCCCACGGC
>$C_2$ (recoded) CTCACTCCTGAGCAGGTCGTCGCAATCGCATCACATGACGGCGGGAAGCAAGCATTGGAGACCGTTCAACGCCTGCTGCCAGTGTTGTGTCAAGCACACGGA
>$T_2$ (original) TTGACCCCCCAGCAGGTGGTGGCCATCGCCAGCAATGGCGGTGGCAAGCAGGCGCTGGAGACGGTCCAGCGGCTGTTGCCGGTGCTGTGCCAGGCCCACGGC
>$T_2$ (recoded) CTGACCCCCCAGCAAGTCGTCGCAATCGCTAGCAATGGTGGAGGTAAACAGGCTCTGGAAACAGTGCAAAGACTTCTCCCCGTCCTCTGCCAAGCCCACGGT
>$G_3$ (original) TTGACCCCCCAGCAGGTGGTGGCCATCGCCAGCAATAATGGTGGCAAGCAGGCGCTGGAGACGGTCCAGCGGCTGTTGCCGGTGCTGTGCCAGGCCCACGGC
>$G_3$ (recoded) TTGACACCTCAACAAGTTGTTGCAATTGCAAGCAATAACGGCGGCAAACAGGCCCTGGAAACTGTGCAGCGCCTTCTCCCAGTTTTGTGTCAGGCACATGGA
>$C_3$ (original) TTGACCCCGGAGCAGGTGGTGGCCATCGCCAGCCACGATGGCGGCAAGCAGGCGCTGGAGACGGTCCAGCGGCTGTTGCCGGTGCTGTGCCAGGCCCACGGC
>$C_3$ (recoded) CTTACTCCAGAACAGGTGGTCGCTATTGCTAGTCATGACGGAGGAAAGCAAGCTCTTGAAACAGTTCAACGGCTCCTTCCTGTTCTTTGCCAAGCTCATGGG
>$T_3$ (original) TTGACCCCCCAGCAGGTGGTGGCCATCGCCAGCAATGGCGGTGGCAAGCAGGCGCTGGAGACGGTCCAGCGGCTGTTGCCGGTGCTGTGCCAGGCCCACGGC
>$T_3$ (recoded) CTCACGCCTCAGCAAGTGGTCGCCATAGCCTCAAATGGAGGAGGTAAGCAAGCATTGGAAACCGTCCAAAGATTGTTGCCCGTGCTCTGCCAGGCCCATGGT
>$G_4$ (original) TTGACCCCCCAGCAGGTGGTGGCCATCGCCAGCAATAATGGTGGCAAGCAGGCGCTGGAGACGGTCCAGCGGCTGTTGCCGGTGCTGTGCCAGGCCCACGGC
>$G_4$ (recoded) CTGACACCTCAACAAGTCGTCGCAATTGCTTCCAACAATGGAGGCAAGCAGGCTCTGGAAACTGTCCAGAGGCTGCTCCCCGTCTTGTGTCAAGCTCATGGT
>$C_4$ (original) TTGACCCCGGAGCAGGTGGTGGCCATCGCCAGCCACGATGGCGGCAAGCAGGCGCTGGAGACGGTCCAGCGGCTGTTGCCGGTGCTGTGCCAGGCCCACGGC
>$C_4$ (recoded) TTGACTCCAGAGCAGGTTGTTGCTATAGCATCACATGACGGCGGAAAACAGGCCCTGGAGACCGTGCAGAGACTTCTTCCTGTGCTTTGCCAAGCACACGGC
>$T_4$ (original) TTGACCCCCCAGCAGGTGGTGGCCATCGCCAGCAATGGCGGTGGCAAGCAGGCGCTGGAGACGGTCCAGCGGCTGTTGCCGGTGCTGTGCCAGGCCCACGGC
>$T_4$ (recoded) CTCACCCCTCAACAAGTAGTGGCCATTGCCTCTAACGGGGGCGGTAAGCAAGCATTGGAAACAGTTCAACGCTTGCTGCCAGTCCTCTGTCAGGCCCATGGG
>$G_5$ (original) TTGACCCCCCAGCAGGTGGTGGCCATCGCCAGCAATAATGGTGGCAAGCAGGCGCTGGAGACGGTCCAGCGGCTGTTGCCGGTGCTGTGCCAGGCCCACGGC
>$G_5$ (recoded) CTCACTCCTCAACAAGTGGTTGCTATTGCTTCTAATAATGGAGGGAAACAAGCTCTTGAAACCGTTCAGCGGCTCCTTCCAGTGCTCTGTCAAGCACATGGA
>$C_5$ (original) TTGACCCCGGAGCAGGTGGTGGCCATCGCCAGCCACGATGGCGGCAAGCAGGCGCTGGAGACGGTCCAGCGGCTGTTGCCGGTGCTGTGCCAGGCCCACGGC
>$C_5$ (recoded) CTTACACCAGAACAGGTTGTCGCAATAGCAAGTCATGACGGGGGCAAGCAGGCCCTTGAGACAGTCCAAAGACTTTTGCCTGTTCTTTGCCAAGCTCACGGC
>$T_5$ (original) TTGACCCCCCAGCAGGTGGTGGCCATCGCCAGCAATGGCGGTGGCAAGCAGGCGCTGGAGACGGTCCAGCGGCTGTTGCCGGTGCTGTGCCAGGCCCACGGC
>$T_5$ (recoded) TTGACCCCCCAACAGGTTGTGGCGATTGCCAGCAATGGGGGGGAAAACAGGCACTGGAAACTGTGCAGAGGTTGCTGCCCGTCTTGTGCCAGGCACACGGG
>half-T (original) TTGACCCCTCAGCAGGTGGTGGCCATCGCCAGCAATGGCGGCGGCAGGCCGGCGCTGGAG
>half-T (recoded) TTAACGCCCCAACAGGTCGTTGCGATCGCGTCTAATGGAGGGGGCAGACCAGCGTTGGAG

Right TALEN arm Binding domain: $G_1T_1G_2A_1T_2C_1C_2C_3C_4C_5C_6A_2G_3C_7A_3$

| | |
|---|---|
| >$G_1$ (original) | TTGACCCCCAGCAGGTGGTGGCCATCGCCAGCAATAATGGTGGCAAGCAGGCGCTGGAGACGGTCCAGCGGCTGTTGCCGGTGCTGTGCCAGGCCCACGGC |
| >$G_1$ (recoded) | CTCACACCTCAACAGGTTGTTGCCATTGCATCAAACAATGGAGGTAAACAAGCACTGGAAACCGTTCAGAGACTTCTCCCTGTCCTTTGTCAGGCTCATGGT |
| >$T_1$ (original) | TTGACCCCCAGCAGGTGGTGGCCATCGCCAGCAATGGCGGTGGCAAGCAGGCGCTGGAGACGGTCCAGCGGCTGTTGCCGGTGCTGTGCCAGGCCCACGGC |
| >$T_1$ (recoded) | CTGACTCCCCAGCAGGTCGTGGCTATCGCCTCTAATGGGGGCGGAAAGCAGGCTTTGGAGACAGTGCAAAGGCTGCTGCCCGTTCTCTGCCAGGCCCACGGG |
| >$G_2$ (original) | TTGACCCCCAGCAGGTGGTGGCCATCGCCAGCAATAATGGTGGCAAGCAGGCGCTGGAGACGGTCCAGCGGCTGTTGCCGGTGCTGTGCCAGGCCCACGGC |
| >$G_2$ (recoded) | CTTACACCACAACAGGTGGTTGCTATTGCTAGTAACAACGGCGGAAAACAAGCCCTCGAAACTGTCCAGAGGCTCTTGCCTGTCCTTTGTCAAGCCCATGGT |
| >$A_1$ (original) | TTGACCCCGGAGCAGGTGGTGGCCATCGCCAGCAATATTGGTGGCAAGCAGGCGCTGGAGACGGTGCAGGCGCTGTTGCCGGTGCTGTGCCAGGCCCACGGC |
| >$A_1$ (recoded) | TTGACCCCTGAACAAGTGGTAGCGATTGCGAGCAACATCGGGGGGAAGCAAGCACTGGAGACTGTGCAAGCCCTGCTCCCAGTGCTGTGTCAGGCTCATGGC |
| >$T_2$ (original) | TTGACCCCCAGCAGGTGGTGGCCATCGCCAGCAATGGCGGTGGCAAGCAGGCGCTGGAGACGGTCCAGCGGCTGTTGCCGGTGCTGTGCCAGGCCCACGGC |
| >$T_2$ (recoded) | CTGACCCCCCAGCAAGTCGTGGCAATCGCCTCCAATGGCGGAGGTAAACAGGCTCTTGAAACAGTCCAGAGACTTCTGCCCGTCCTCTGCCAAGCCCACGGT |
| >$C_1$ (original) | TTGACCCCGGAGCAGGTGGTGGCCATCGCCAGCCACGATGGCGGCAAGCAGGCGCTGGAGACGGTCCAGCGGCTGTTGCCGGTGCTGTGCCAGGCCCACGGC |
| >$C_1$ (recoded) | CTTACTCCAGAACAAGTCGTCGCCATCGCTAGCCACGACGGGGGCAAACAGGCCCTGGAGACTGTGCAACGCTTGCTTCCAGTTTTGTGCCAGGCACATGGA |
| >$C_2$ (original) | TTGACCCCGGAGCAGGTGGTGGCCATCGCCAGCCACGATGGCGGCAAGCAGGCGCTGGAGACGGTCCAGCGGCTGTTGCCGGTGCTGTGCCAGGCCCACGGC |
| >$C_2$ (recoded) | CTCACTCCTGAGCAGGTCGTGGCAATCGCATCACATGATGGTGGGAAGCAAGCATTGGAGACCGTTCAGCGCTTGCTGCCAGTGCTGTGTCAAGCACACGGA |
| >$C_3$ (original) | TTGACCCCGGAGCAGGTGGTGGCCATCGCCAGCCACGATGGCGGCAAGCAGGCGCTGGAGACGGTCCAGCGGCTGTTGCCGGTGCTGTGCCAGGCCCACGGC |
| >$C_3$ (recoded) | TTGACCCCGAACAGGTCGTCGCCATTGCTAGTCACGACGGAGGAAAGCAGGCTCTTGAAACAGTTCAAAGGCTCCTTCCTGTCCTTTGCCAGGCTCATGGG |
| >$C_4$ (original) | TTGACCCCGGAGCAGGTGGTGGCCATCGCCAGCCACGATGGCGGCAAGCAGGCGCTGGAGACGGTCCAGCGGCTGTTGCCGGTGCTGTGCCAGGCCCACGGC |
| >$C_4$ (recoded) | CTGACTCCAGAGCAAGTTGTTGCTATCGCCTCACATGATGGGGGAAAACAGGCCCTCGAGACCGTGCAGAGACTGTTGCCTGTTCTCTGCCAAGCACACGGA |
| >$C_5$ (original) | TTGACCCCGGAGCAGGTGGTGGCCATCGCCAGCCACGATGGCGGCAAGCAGGCGCTGGAGACGGTCCAGCGGCTGTTGCCGGTGCTGTGCCAGGCCCACGGC |
| >$C_5$ (recoded) | CTTACACCCGAACAGGTGGTCGCAATAGCAAGCCACGACGGTGGTAAGCAGGCACTTGAGACAGTCCAAAGACTTCTCCCTGTGCTTTGCCAGGCCCACGGT |
| >$C_6$ (original) | TTGACCCCGGAGCAGGTGGTGGCCATCGCCAGCCACGATGGCGGCAAGCAGGCGCTGGAGACGGTCCAGCGGCTGTTGCCGGTGCTGTGCCAGGCCCACGGC |
| >$C_6$ (recoded) | CTCACTCCAGAGCAGGTTGTTGCCATTGCCTCCCATGATGGAGGGAAACAAGCTCTCGAAACTGTTCAGAGGTTGCTTCCCGTCCTCTGTCAAGCTCACGGG |
| >$A_2$ (original) | TTGACCCCGGAGCAGGTGGTGGCCATCGCCAGCAATATTGGTGGCAAGCAGGCGCTGGAGACGGTGCAGGCGCTGTTGCCGGTGCTGTGCCAGGCCCACGGC |
| >$A_2$ (recoded) | CTGACGCCAGAGCAAGTGGTGGCCATCGCTTCTAATATCGGCGGCAAGCAGGCCCTGGAAACAGTTCAAGCTTTGCTACCCGTGTTATGCCAGGCCCACGGC |
| >$G_3$ (original) | TTGACCCCCAGCAGGTGGTGGCCATCGCCAGCAATAATGGTGGCAAGCAGGCGCTGGAGACGGTCCAGCGGCTGTTGCCGGTGCTGTGCCAGGCCCACGGC |
| >$G_3$ (recoded) | CTGACCCCTCAGCAGGTTGTGGCAATTGCATCCAACAACGGTGGGAAACAAGCACTCGAAACTGTGCAGCGCCTTCTCCCAGTTTTGTGTCAAGCACATGGA |
| >$C_7$ (original) | TTGACCCCGGAGCAGGTGGTGGCCATCGCCAGCCACGATGGCGGCAAGCAGGCGCTGGAGACGGTCCAGCGGCTGTTGCCGGTGCTGTGCCAGGCCCACGGC |
| >$C_7$ (recoded) | TTGACACCCGAACAGGTTGTCGCTATCGCCAGTCATGACGGCGGTAAGCAGGCCTTGGAGACCGTCCAGAGGCTCCTTCCCGTGCTGTGCCAGGCACATGGG |
| >$A_3$ (original) | TTGACCCCGGAGCAGGTGGTGGCCATCGCCAGCAATATTGGTGGCAAGCAGGCGCTGGAGACGGTGCAGGCGCTGTTGCCGGTGCTGTGCCAGGCCCACGGC |
| >$A_3$ (recoded) | CTAACCCCGGAGCAGGTAGTGGCCATTGCTTCTAATATTGGGGGCAAACAAGCTCTCGAGACGGTACAGGCCCTCTTGCCAGTTTTGTGTCAAGCGCACGGC |
| >half-T (original) | TTGACCCCTCAGCAGGTGGTGGCCATCGCCAGCAATGGCGGCGGCAGGCCGGCGCTGGAG |
| >half-T (recoded) | TTAACCCCCCAACAGGTCGTTGCGATCGCGTCTAATGGAGGGGGCAGACCAGCGTTGGAG |

Left TALEN arm (Identity between motifs: 94.1%-100%)

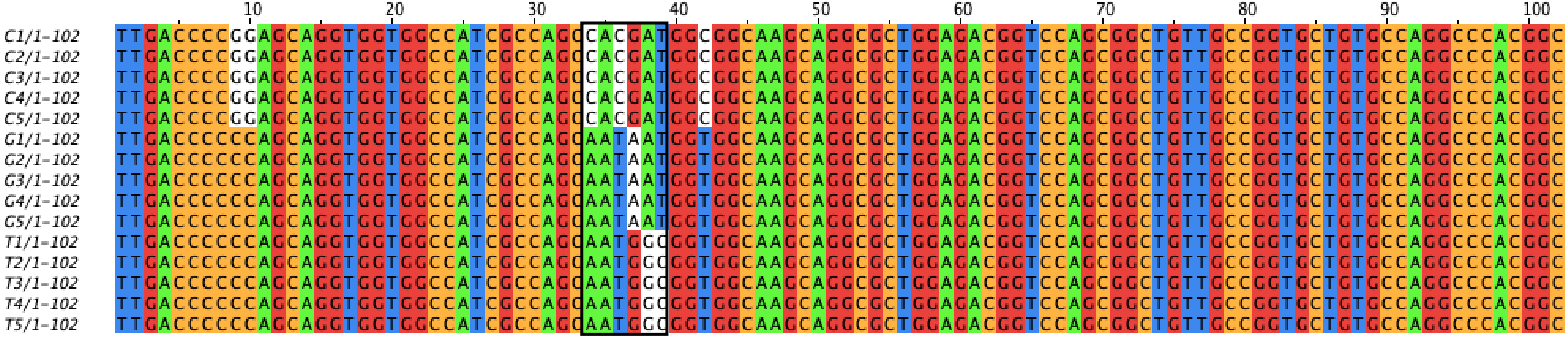


Right TALEN arm (Identity between motifs: 94.1%-100%)

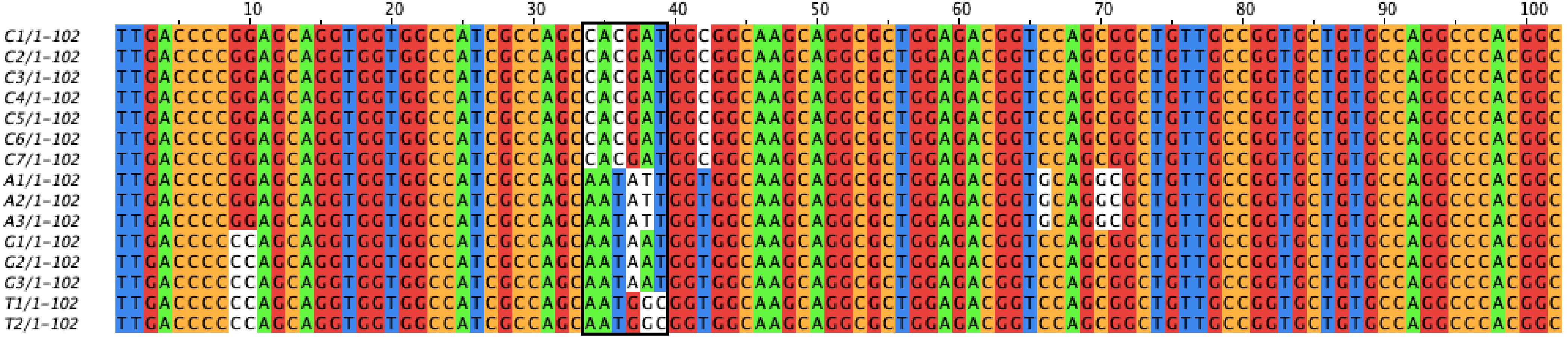

Left $TALEN_{RECODED}$ arm (Identity between motifs: 66.7%-78.4%)

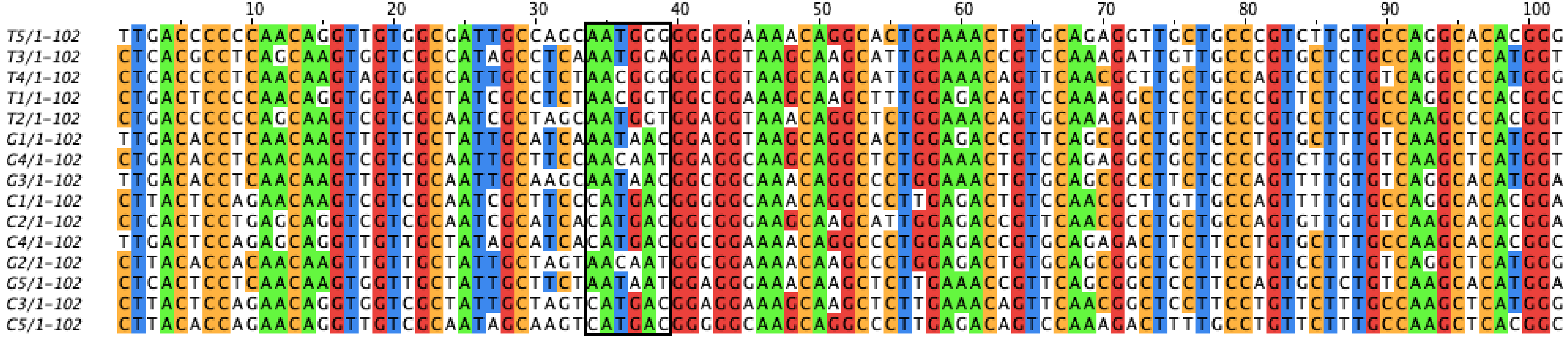


Right $TALEN_{RECODED}$ arm (Identity between motifs: 65.7%-78.4%)

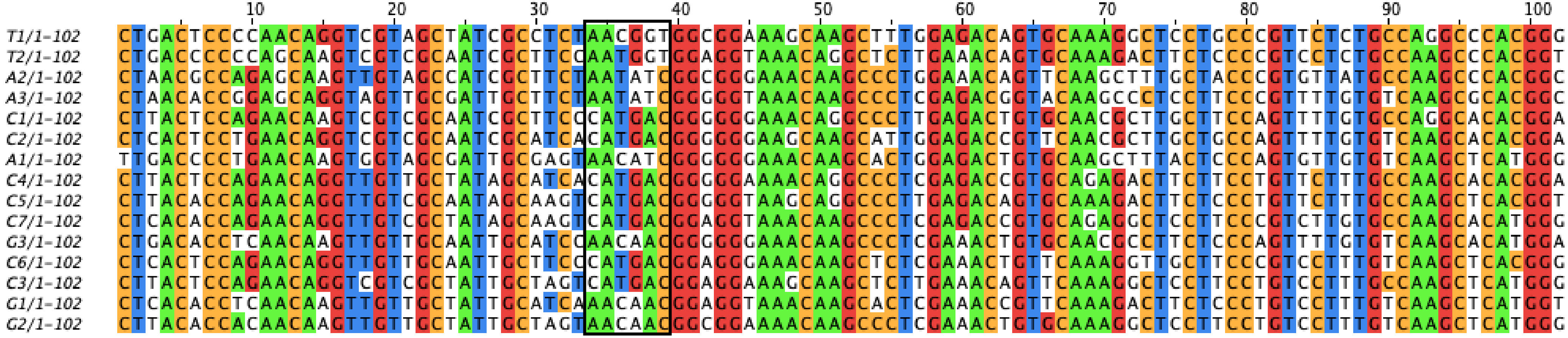
