## Supplemental Table S1 for "TALEN-induced contraction of CTG trinucleotide repeats in myotonic dystrophy type 1 cells"

**Supplemental Table S1: PacBio sequencing data of the DM1 reference genome (ASA3-p11)**

| | |
|---|---|
| Nucleotides covered by subreads: | 1381 Gb |
| Nucleotides covered by HiFi reads: | 82.5 Gb |
| Total coverage: | 13 X |

**Assembly (primary)**

| | | |
|---|---|---|
| Contigs | (total) | 371 |
| | > 1 kb | 371 |
| | > 5 kb | 371 |
| | > 10 kb | 369 |
| | > 25 kb | 331 |
| | > 50 kb | 291 |
| Largest contig | | 160.3 Mb |
| Total length | | 3.044 Gb |
| GC (%) | | 40.8 |
| N50 | | 52.9 Mb |
| N90 | | 9.2 Mb |
| L50 | | 17 |
| L90 | | 69 |
| N's per 100 kb | | 0.00 |
