## Supplemental Table S2 for "TALEN-induced contraction of CTG trinucleotide repeats in myotonic dystrophy type 1 cells"

**Supplemental Table S2: Summary of PacBio sequencing**

| PacBio Sequel | Clone | TALEN arms | Nbr SMRTCells |
|---|---|---|---|
| | ASA03-44 | Right+Left | 4 |
| | ASA2 | Right+Left | 4 |
| | ASA22C | Right+Left | 4 |
| | ASA27 | Right+Left | 4 |
| | ASA28 | Right+Left | 4 |
| | ASA30 | Right+Left | 4 |
| | ASA51 | Right+Left | 4 |
| | ASA53 | Right+Left | 4 |
| | ASA54 | Right+Left | 2 |
| | ASA111E | Right+Left | 4 |
| | **Total** | | **38** |

| PacBio Sequel II | Clone | TALEN arms | Nbr SMRTCells |
|---|---|---|---|
| (Reference genome) | ASA3-p11 | None | 4 |
| | ASA03-44 | Right | 3 |
| | ASA27 | Right+Left | 3 |
| | ASA53 | Right+Left | 3 |
| | ASA54 | Right+Left | 1 |
| | C3_gauche | Left recoded | 3 |
| | C5_gauche | Left recoded | 3 |
| | C3_TALEN | (Right+Left)recoded | 2 |
| | C6_TALEN | (Right+Left)recoded | 2 |
| | C9_TALEN | (Right+Left)recoded | 3 |
| | C11_TALEN | (Right+Left)recoded | 3 |
| | C12_TALEN | (Right+Left)recoded | 2 |
| | C15_TALEN | (Right+Left)recoded | 2 |
| | C16_TALEN | (Right+Left)recoded | 1 |
| | **Total** | | **35** |
