## Supplemental Table S3 for "TALEN-induced contraction of CTG trinucleotide repeats in myotonic dystrophy type 1 cells"

**Supplemental Table S3: Genome coverage**

| Clone | Sequence (nt) | Genome coverage | Genome covered (%) |
|---|---|---|---|
| ASA3-p11 | 8,25E+10 | 13 | 100% |
| ASA03-44 | 3,65E+10 | 6 | 99,7% |
| ASA2 | 2,23E+10 | 4 | 97,3% |
| ASA22C | 2,90E+10 | 5 | 99,1% |
| ASA27 | 7,60E+10 | 12 | 100% |
| ASA28 | 2,22E+10 | 4 | 97,2% |
| ASA30 | 2,08E+10 | 3 | 96,5% |
| ASA51 | 2,36E+10 | 4 | 97,8% |
| ASA53 | 6,21E+10 | 10 | 100% |
| ASA54 | 2,33E+10 | 4 | 97,7% |
| ASA111E | 2,44E+10 | 4 | 98,0% |
| C3_gauche | 1,45E+11 | 23 | 100% |
| C5_gauche | 8,32E+10 | 13 | 100% |
| C3_TALEN | 1,50E+11 | 24 | 100% |
| C6_TALEN | 1,43E+11 | 23 | 100% |
| C9_TALEN | 8,76E+10 | 14 | 100% |
| C11_TALEN | 1,43E+11 | 23 | 100% |
| C12_TALEN | 1,26E+11 | 20 | 100% |
| C15_TALEN | 1,03E+11 | 17 | 100% |
| C16_TALEN | 3,16E+10 | 5 | 99,4% |
